# When to be Temperate: On the Fitness Benefits of Lysis vs. Lysogeny

**DOI:** 10.1101/709758

**Authors:** Guanlin Li, Michael H. Cortez, Jonathan Dushoff, Joshua S. Weitz

**Author notes:** Electronic address; URL: http://ecotheory.biology.gatech.edu.

## Abstract

Bacterial viruses, i.e., ‘bacteriophage’ or ‘phage’, can infect and lyse their bacterial hosts, releasing new viral progeny. In addition to the lytic pathway, certain bacteriophage (i.e., ‘temperate’ bacteriophage) can also initiate lysogeny, a latent mode of infection in which the viral genome is integrated into and replicated with the bacterial chromosome. Subsequently, the integrated viral genome, i.e., the ‘prophage’, can induce and restart the lytic pathway. Here, we explore the relationship between infection mode, ecological context, and viral fitness, in essence asking: when should viruses be temperate? To do so, we use network loop analysis to quantify fitness in terms of network paths through the life history of an infectious pathogen that start and end with infected cells. This analysis reveals that temperate strategies, particularly those with direct benefits to cellular fitness, should be favored at low host abundances. This finding applies to a spectrum of mechanistic models of phage-bacteria dynamics spanning both explicit and implicit representations of intracellular infection dynamics. However, the same analysis reveals that temperate strategies, in and of themselves, do not provide an advantage when infection imposes a cost to cellular fitness. Hence, we use evolutionary invasion analysis to explore when temperate phage can invade microbial communities with circulating lytic phage. We find that lytic phage can drive down niche competition amongst microbial cells, facilitating the subsequent invasion of latent strategies that increase cellular resistance and/or immunity to infection by lytic viruses – notably this finding holds even when the prophage comes at a direct fitness cost to cellular reproduction. Altogether, our analysis identifies broad ecological conditions that favor latency and provide a principled framework for exploring the impacts of ecological context on both the short- and long-term benefits of being temperate.

## I. INTRODUCTION

Viruses of microbes are ubiquitous in natural systems, e.g., densities of virus particles typically exceed 10^7^ per ml in marine systems and 10^8^ per g in soils. Viral infections can transform the fate of target cells, populations, and associated ecosystems (1, 2, 3, 4, 5). Bacteriophage infections can lead to lysis and death of the infected cell, and new infections by progeny virus particles can drive down microbial populations leading to endogenous oscillations in population densities (6, 7). However for many bacteriophage, lysis is not the only possible infection outcome.

Infection by temperate bacteriophage such as phage *λ*, *µ*, and P22 can lead to cell lysis or lysogeny (8, 9). The ‘decision’ process associated with lysis and lysogeny has been termed a genetic switch (10). For example, in phage *λ*, the switch is modulated by a bidirectional promoter that controls expression of regulatory proteins whose stochastic expression and feedback culminates in either lysis or lysogeny (e.g., refer to (11) for a recent review). Notably, the probability of initiating lysogeny and the rate of spontaneous induction are both evolvable traits. For context, induction denotes the excision of the prophage from the bacterial genome and the initiation of the lytic pathway. The evolvability of quantitative traits associated with temperate phage raises the question: how do the benefits of lysogeny vary with ecological conditions?

Prior hypotheses had suggested that temperate phage have an evolutionary advantage when few hosts are available and extracellular virion decay rates are high (12). In 1984, Frank Stewart and Bruce Levin addressed this question by analyzing nonlinear dynamics models of nutrients, cells, virulent phage and temperate phage (13). However, Stewart and Levin reported that “in spite of the intuitive appeal of this low density hypothesis, we are unable to obtain solutions consistent with it using the model presented here.” Instead, they introduced external oscillations in resource supply rates in order to identify regimes in which both virulent and temperate bacteriophage could coexist. Moreover, by analyzing population abundances as a proxy for evolutionary success, the ‘advantage’ of a temperate vs. obligately lytic strategy was compared in terms of relative abundances of virus particles and infected cells. Such weightings are seemingly arbitrary and do not stem from an evolutionary framework.

Here, we re-assess the benefits of being temperate by asking the question: under what ecological conditions can temperate phage potentially invade microbial communities, including those without and with circulating lytic phage? First, we adapt a cell-centric metric of viral invasion fitness to the ecological dynamics of viruses and their microbial hosts (14–16). Using a novel application of Levins’ network loop analysis (17), we show that complicated algebraic expressions for viral fitness can be biologically interpreted in terms of purely horizontal, vertical, and mixed transmission pathways. In doing so, we show that temperate strategies can invade virus-free environments when susceptible densities are relatively low and when the integrated prophage confers direct fitness benefits to cellular growth and survival. However, prophage can sometimes impose a cost to cellular growth and survival. Hence, we use evolutionary invasion analysis to identify when temperate phage can successfully invade a community including bacteria and circulating lytic phage. As we show, lytic phage can drive down microbial cell densities so as to enable invasion by temperate phage that confer protection against subsequent infection. This result holds even when the temperate phage provide no direct benefit or even impose a cost to cellular growth. Overall, this analysis provides a theoretical framework for identifying near-term ‘solutions’ (sensu Stewart and Levin) to the problem of when to be temperate in an ecological context. As we discuss, identifying such near-term solutions also opens the door to new approaches to addressing the long-term evolution of temperate strategies.

## II. RESULTS

### A. Nonlinear, Population Model of Temperate Phage Dynamics

We begin by considering the dynamics of temperate phage in a nonlinear population model that includes explicit representation of infections including cells that are either susceptible (*S*), exposed (*E*), actively infected (*I*), or lysogens (*L*), as well as virus particles (*V*), see the top panel of Figure 1.The exposed cells are cells that have been infected but the virus has not yet committed to either the lytic pathway (turning it into an actively infected, *I*, cell) or the lysogenic pathway (turning it into a lysogen, *L*). In this model, the life history traits of a temperate phage are defined by two evolvable parameters: *p*, the probability a virus enters the lysogenic pathway and *γ*, the induction rate after a virus enters the lysogenic pathway. We represent this model in terms of a system of nonlinear ordinary differential equations (ODEs):

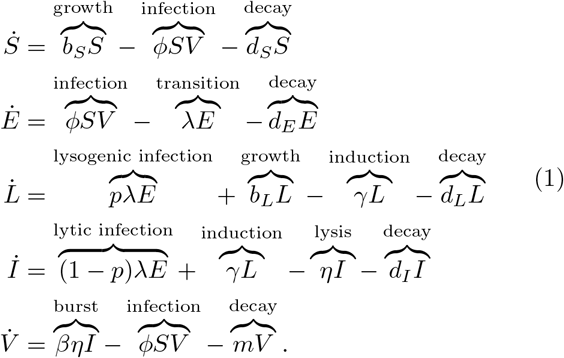

**FIG. 1:**
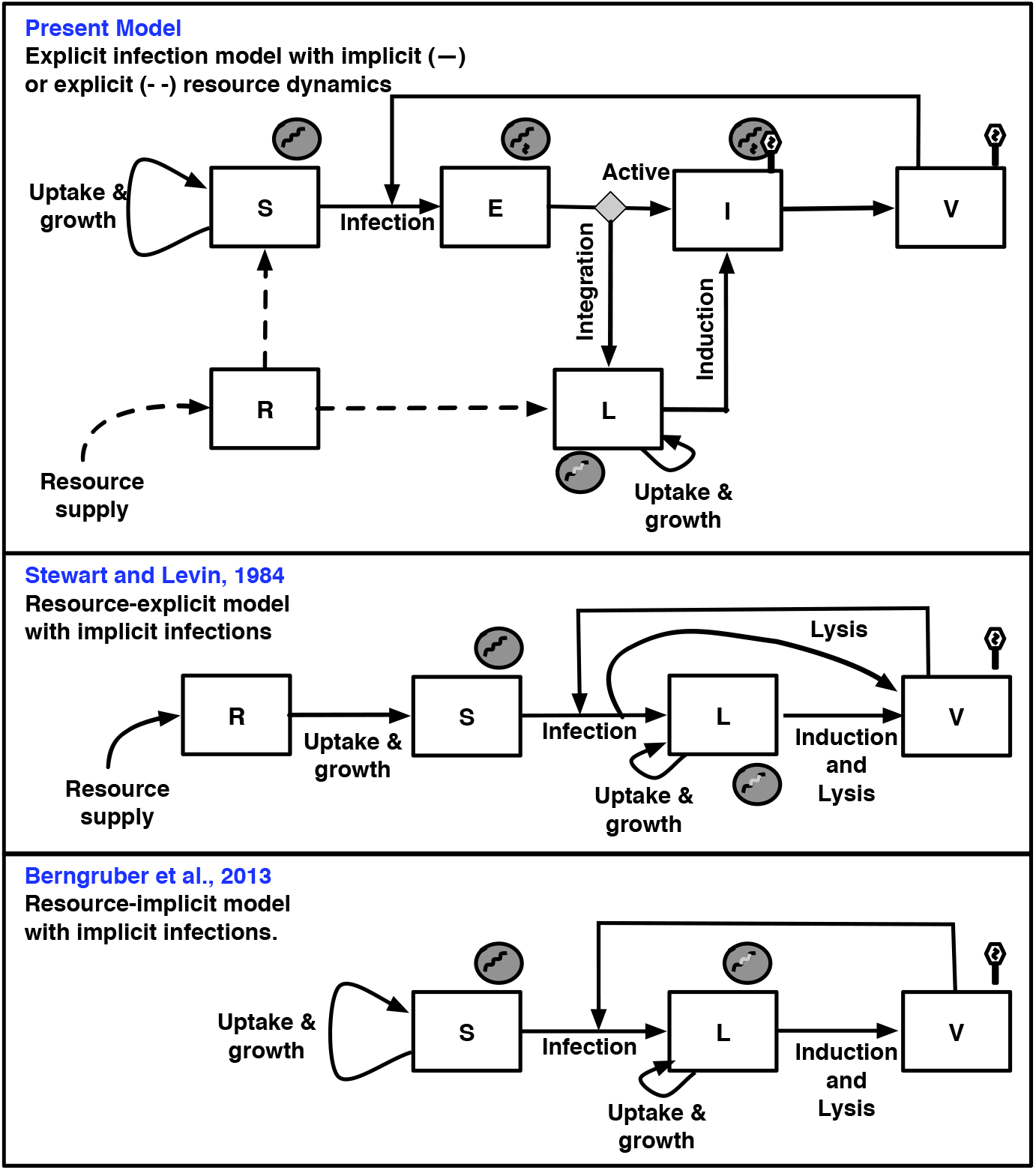
Schematic of nonlinear dynamics population models of temperate phage. (Top) The explicit infection model with implicit (–) or explicit (- -) resource dynamics (developed here). (Middle) Resource-explicit model with implicit infections (13). (Bottom) Resource-implicit model with implicit infections (15). The governing equations for each model are presented in the Main Text (Top), Methods (Middle and Bottom), with extended analysis in Appendix A.

In this model, *φ* is the adsorption rate, *d*_*S*_, *d*_*E*_, *d*_*L*_ and *d*_*I*_ are the cellular death rates of susceptible cells, exposed infected cells, lysogens and lytic-fated infected cells respectively, *λ* is the transition rate from exposed cells to the fate determined cells, *p* is the probability of lysogeny, *γ* is the induction rate, *η* is the lysis rate, *β* is the burst size and *m* is the virion decay rate. The growth rates of susceptible hosts and lysogens are denoted by *b*_*S*_ and *b*_*L*_, respectively. For the resource-implicit model, the growth rates are

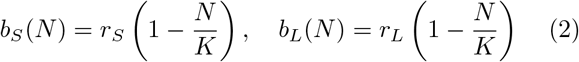

where *N* = *S* + *E* + *L* + *I* is the population density of total cells. Parameters *r*_*S*_ and *r*_*L*_ denote the maximal cellular growth rates of susceptible cells and lysogens, *K* is the carrying capacity. For the resource-explicit model, the growth rates of susceptible hosts and lysogens are

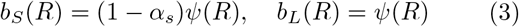

where *R* is the resource density and *ψ*(*R*) = *µ*_*max*_*R*/(*R*_*in*_ + *R*) is the Monod equation. For the resource-explicit model, we add one additional equation to describe the dynamics of the resources,

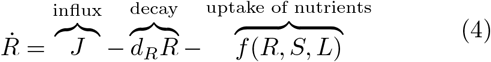

where *f* (*R, S, L*) = *eψ*(*R*) (*L* + (1 − *α*_*s*_)*S*) denotes the cumulative uptake of nutrients by all cells. Parameters *µ*_*max*_ and *R*_*in*_ are the maximal cellular growth rate and the half-saturation constant, *J* and *d*_*R*_ are the influx and decay rates of resources, *e* is the conversion efficiency, *α*_*s*_ is the selection coefficient that measures the relative difference in the reproductive output between lysogens and susceptible cells. Model 1 allows us to analyze the dynamics of viruses with different life history strategies. Here, a viral strategy is defined by different combinations of the trait values (*p*, *γ*). In the twodimensional viral strategy space we denote the purely lytic strategy as *p* = 0 (for which the value of *γ* is irrelevant, and assumed to be *γ*_*max*_ for convenience), and the purely lysogenic strategy as *p* = 1, *γ* = *γ*_*min*_, where *γ*_*min*_ > 0. Here, a temperate viral strategy is characterized by 0 *< p* ≤ 1 and *γ* ≥ *γ*_*min*_. This model extends earlier proposals to model temperate phage via implicit infections (i.e., without an explicit state that demarcates the decision between lysis and lysogeny), either with explicit resource dynamics (13) or with implicit resource dynamics (15). Critically, all of these models include the possibility of both vertical and horizontal transmission of phage genomes. These models also set the basis for evaluating how temperate phage may invade in a disease-free context, i.e., given the equilibrium concentrations of *S*^∗^ in resource-implicit cases or (*R*^∗^, *S*^∗^) in resource-explicit cases.

### B. Viral invasion analysis

Here, we show how our model can be used to compute viral invasion fitness for viruses with different lysogeny-lysis strategies, *i.e.*, combinations of trait values (*p*, *γ*). The spread of temperate viruses in a parasite-free environment can be analyzed in terms of the basic reproduction number 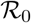, which denotes the average number of new infected cells produced by a single (typical) infected cell and its progeny virions in an otherwise susceptible population (14, 15, 16). When 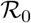 is greater than 1, the pathogen will spread and when 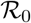 is less than 1, the pathogen will not spread. The next-generation matrix (NGM) approach can be used to calculate 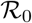 (18). The NGM represents the expected progeny for transitions between all combinations of ‘epidemiological birth states’, *i.e.*, states that can produce newly infected hosts cells. In our model, states *E* and *L* are the only epidemiological birth states and epidemiological births arise due to infection of cells by virions and by the division of lysogenic cells. The largest positive eigenvalue of the NGM is equivalent to 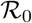. The NGM for model 1 is the 2 *×* 2 matrix Φ:

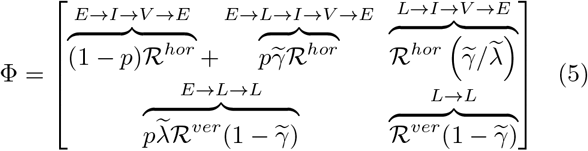

Here 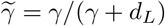 is the probability induction occurs before cell death, 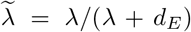 is the probability that exposed cells enter the lysogenic pathway before cell death, 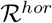 and 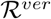 are the basic reproduction numbers of purely lytic phage (*p* = 0) and purely lysogenic phage (*p* = 1, *γ* = 0), respectively:

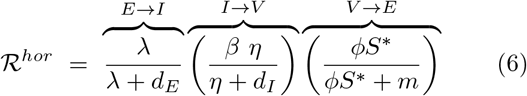

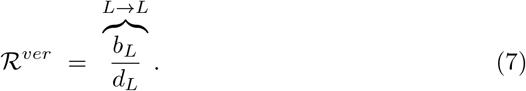

where *S*^∗^ = *K*(1 − *d*_*S*_/*r*_*S*_) is the susceptible host density in the virus-free environment. The vertical contribution to fitness is modulated by infected cell growth rates, *b*_*L*_, which in the resource-implicit and resource-explicit systems are *r*_*L*_ (1 − *S*^∗^/*K*) and *ψ*(*R*^∗^) respectively, where *R*^∗^ is the resource concentration in the virus-free environment.

Each entry Φ_*ij*_ in the NGM represents the expected number of new infected individuals in epidemiological birth state *i* (either *L* or *E*), generated by one infected individual at epidemiological birth state *j* (either *L* or *E*), accounting for new infections that arise via the lytic and lysogenic pathways. For example, Φ_11_ accounts for the expected number of exposed cells *E* produced by a single exposed cell. The single exposed cell *E* has a probability 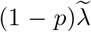 of entering the *I* state, of which a fraction *η*/(*η* + *d*_*I*_) of infected cells will release viruses. Each infected cell produces *β* free virus particles given successful lysis. The factor *φS*^∗^/(*φS*^∗^ + *m*) denotes the probability that a free virus particle is adsorbed into a susceptible cell before it decays. As such, 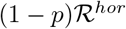 is the expected number of newly infected exposed cells produced by a single exposed cell via a purely lytic pathway. Similarly, an exposed cell *E* has a probability 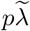 of entering the *L* state and being induced from the lysogenic cell *L* to infected cell *I*. Thus, a single exposed cell produces 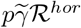 newly infected exposed cells on average after a sequence of integration, induction, and lysis events. Altogether, the expected number of exposed cells produced by a single exposed cell is 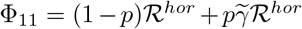. The other entries in the NGM can be interpreted similarly; the transmission pathways are labeled in Eq. [5] for each entry of the NGM.

In the event that phage cannot induce, i.e., *γ* = 0, then the basic reproduction number reduces to:

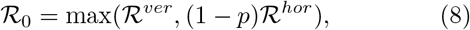

which corresponds to the viral invasion fitness associated with either lysis or lysogeny, but not both, equivalent to the finding in (16).

The use of a cell-centric metric enables direct comparisons of lytic and lysogenic pathways, i.e., even in the absence of virion production from lysogens. In the general case where induction is possible, i.e., *γ >* 0, then, the basic reproduction number 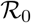 becomes:

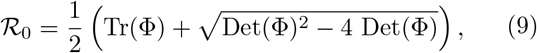

where the trace Tr(Φ) is:

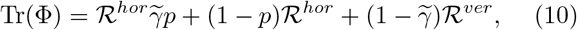

and the determinant Det(Φ) is:

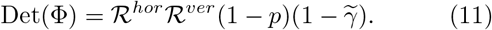

Notably, this formula for the basic reproduction number applies to all model variants of temperate phage dynamics listed in Figure 1, see Appendix B2 for details. However, Eq. [9] poses multiple challenges for interpreting 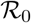 not only for phage, but for generalized cases of host-pathogen dynamics with multiple transmission modes (19). It is the interpretation problem that we address next.

### C. Interpreting 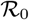 using Levins’ loop analysis

Here, we use Levins’ loop analysis (17) to interpret the basic reproduction number 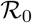 arising in each of the models depicted in Figure 1. Levins’ loop analysis was developed for the analysis of feedback in ecological networks (17) - which we adapt to the study of the network of paths in the life history of an infectious pathogen. Loop analysis has been used previously to interpret 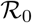 for discrete-time stage structured population dynamics (20). In another instance, previous work (21) focused on applications of loop analysis to models where the NGM only has a single non-zero eigenvalue whereas the NGM given by Eq. [5] has multiple non-zero eigenvalues. In the present context, we define a one-generation loop as the collection of paths that start in one infection class and ends in the same class without revisiting any classes. Next, we define a joint two-generation loop as a pair of one-generation loops that start and end in the same class. Finally, we define a disjoint two-generation loop as a pair of one-generation loops whose pathways do not share an epidemiological birth state.

Using these definitions, we denote the lytic loop as ①. In this loop, a newly infected individual in state *E* passes to state *I* and releases virions that lead to new hosts entering the *E* state. The first loop in Figure 2 shows the pathway from *E* to *I* to *V* and then back to *E*. We denote the lysogenic loop as ②. In this loop, lysogens are reproduced during the lifespan of an individual lysogen. The second loop in Figure 2 shows the pathway from *L* to *L*. We denote the lyso-lytic loop as ③. In this loop, a newly infected individual in state *E* passes to then releases virions that lead to new hosts entering the *E* state. The third loop in Figure 2 shows the pathway from *E* to *L* to *I* to *V* and then back to *E*.

**FIG. 2:**
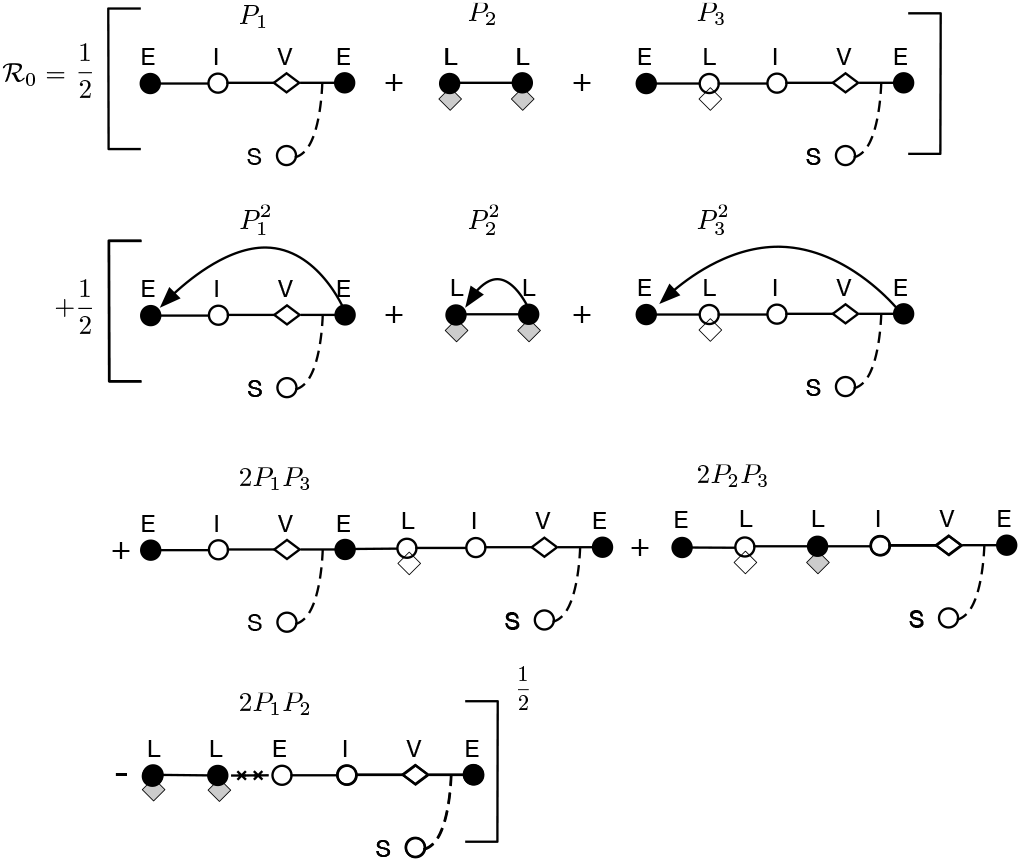
Loop-based interpretation of the basic reproduction number for horizontal and vertical transmission of temperate phage. Closed circles denote epidemiological births, open circles represent infected cell transitions, diamonds represent virus particles, and lysogens are denoted using a hybrid symbol (denoting the presence of an integrated viral genome). Solid lines denote transitions between states, lines with arrows denote a repeat of the same loop, dashed lines denote interactions with uninfected cells, and crossed out lines (-x-x-) denote non-feasible transitions.

We can compute a reproduction number for each loop, *i.e.*, the expected number of new infections that arises from each loop. We denote the reproduction number of the lytic loop ① as *P*_1_, the reproduction number of the lysogenic loop ② as *P*_2_ and the reproduction number of the lyso-lytic loop ③ as *P*_3_. The reproduction numbers of one-generation loops can be directly read-off from the NGM Φ, Eq. [5],

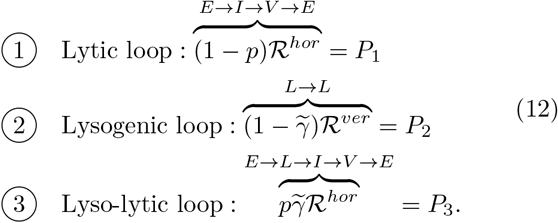

Among all the pairs of one-generation loops, there are seven joint two-generation loops with permutations, e.g. ① ⊕ ①, ② ⊕ ②, ③ ⊕ ③, ① ⊕ ③, ③ ⊕ ①, ② ⊕ ③ and ③ ⊕ ②, and two disjoint pairs of loops, e.g. ① ⊕ ② and ② ⊕ ①. Using the above, Eq. [9] for 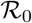 can be rewritten in the following form:

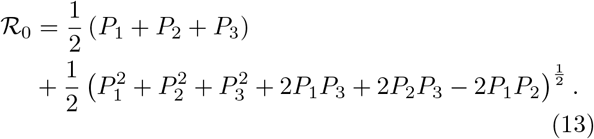

Notice that *P*_1_ + *P*_2_ + *P*_3_ represents the sum of the one-generation loops and the terms in the square root represent the sum of the joint two-generation loops discounted by the disjoint loops. The contributions from all two-generation loops are discounted by the 1*/*2 exponent because the basic reproductive number focuses on reproductive output after one generation. Figure 2 shows the equivalency between Eq. [13] and the loop-based interpretation.

Notably, the calculations of 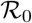 for all temperate phage models in Figure 1 can be expressed in the form of Eq. [13]. Table (I) summarizes the equivalencies of the component calculations for 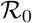 in each of the models given a lytic loop, lysogenic loop, and a mixed loop. Moreover, this loop interpretation also applies to reduced versions of these models as long as they retain two epidemiological birth states (e.g., the *SILV* -system, as analyzed in (16)). Although the models differ in their mechanistic details, each shares two evolvable traits: *p*, the probability of entering lysogenic state after infection and *γ*, the rate of induction from a lysogenic state. This similarity in form suggests that the dependence for invasion on the temperate traits *p* and *γ* may also transcend model details.

**TABLE 1:** Loop-based 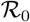 Calculations for Temperate Phage Models

| Temperate phage dynamics | Lytic loop<br>$P_1$ | Lysogenic loop<br>$P_2$ | Lyso-lytic loop<br>$P_3$ |
| --- | --- | --- | --- |
| Resource-explicit model with implicit infections (Stewart & Levin, 1984) | $\overbrace{(1-p) \left( \frac{\beta \phi S^*}{\rho} \right)}^{V \rightarrow V}$ | $\overbrace{\frac{\psi(R^*)}{\gamma + \rho + \nu}}^{L \rightarrow L}$ | $\overbrace{\left( \frac{\gamma}{\gamma + \rho + \nu} \right)}^{L \rightarrow V} \overbrace{\left( \frac{\beta \phi S^*}{\rho} \right)}^{V \rightarrow L} p$ |
| Resource-implicit model with implicit infections (Berngruber, et al., 2013) | $\overbrace{(1-p) \left( \frac{b\beta \phi S^*}{\phi S^* + m} \right)}^{V \rightarrow V}$ | $\overbrace{\frac{r_L \delta (1 - S^*/K)}{m + \gamma}}^{L \rightarrow L}$ | $\overbrace{\left( \frac{\gamma}{\gamma + m} \right)}^{L \rightarrow V} \overbrace{\left( \frac{b\beta \phi S^*}{\phi S^* + m} \right)}^{V \rightarrow L} p$ |
| Resource-explicit model with explicit infections (Present) | $\overbrace{(1-p) \left( \frac{\lambda}{\lambda + d_E} \right)}^{E \rightarrow I} \overbrace{\left( \frac{\beta \eta}{\eta + d_I} \right)}^{I \rightarrow V} \overbrace{\left( \frac{\phi S^*}{\phi S^* + m} \right)}^{V \rightarrow E}$ | $\overbrace{\frac{r_L (1 - S^*/K)}{\gamma + d_L}}^{L \rightarrow L}$ | $p \overbrace{\left( \frac{\lambda}{\lambda + d_E} \right)}^{E \rightarrow L} \overbrace{\left( \frac{\gamma}{\gamma + d_L} \right)}^{L \rightarrow I} \overbrace{\left( \frac{\beta \eta}{\eta + d_I} \right)}^{I \rightarrow V} \overbrace{\left( \frac{\phi S^*}{\phi S^* + m} \right)}^{V \rightarrow E}$ |
| Resource-implicit model with explicit infections (Present) | $\overbrace{(1-p) \left( \frac{\lambda}{\lambda + d_E} \right)}^{E \rightarrow I} \overbrace{\left( \frac{\beta \eta}{\eta + d_I} \right)}^{I \rightarrow V} \overbrace{\left( \frac{\phi S^*}{\phi S^* + m} \right)}^{V \rightarrow E}$ | $\overbrace{\frac{\psi(R^*)}{\gamma + d_L}}^{L \rightarrow L}$ | $p \overbrace{\left( \frac{\lambda}{\lambda + d_E} \right)}^{E \rightarrow L} \overbrace{\left( \frac{\gamma}{\gamma + d_L} \right)}^{L \rightarrow I} \overbrace{\left( \frac{\beta \eta}{\eta + d_I} \right)}^{I \rightarrow V} \overbrace{\left( \frac{\phi S^*}{\phi S^* + m} \right)}^{V \rightarrow E}$ |

### D. Feasible invasion strategies

We systematically analyze the dependency of the viral invasion fitness 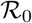 on the viral ‘strategy’, *i.e.*, the combination of temperate traits (*p, γ*), for all four temperate phage models depicted in Figure 1. A strategy (*p, γ*) is feasible if 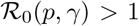. We conduct this invasion analysis to distinguish when a temperate strategy is feasibly and/or obligately invasible. For feasible invasibility, we identify the ecological conditions in which a temperate viral strategy can invade in a completely susceptible host population (*i.e.*, which combination of traits corresponds to 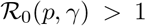. For obligate invasibility, we identify the ecological conditions in which a temperate viral strategy is required for invasion of a completely susceptible host population such that 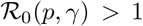 for all 0 *< p ≤* 1 whereas the purely lytic strategy cannot invade, *i.e.*, 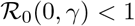.

To begin, it is useful to delineate the bounds to 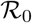 in special cases. As shown in the Appendix C2, we find that in the case of a resource-implicit model the purely lytic strategy maximizes 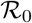 when 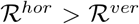, *i.e.*, more newly infected cells are produced through the lytic pathway than the lysogenic pathway. In contrast, the purely lysogenic strategy maximizes 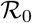 when 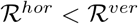, *i.e.*, more newly infected cells are produced through the lysogenic pathway than the lytic pathway. Notably, 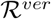 decreases with susceptible host density *S*^∗^ while 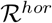 increases with susceptible host density *S*^∗^. Hence, 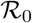 is higher for temperate phage until *S*^∗^ is sufficiently high that 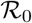 is higher for lytic phage. For the resource-explicit models, the virus-free environment is represented by the susceptible host density (*S*^∗^) and resource density (*R*^∗^). In the *S*^∗^-*R*^∗^ plane, there is a critical transition curve (*S*_*c*_, *R*_*c*_) defined by 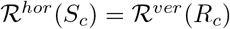 where the strategy associated with maximal fitness (*i.e.*, that maximizes 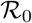) switches from purely lysogenic to purely lytic (see Appendix C2). This analysis reveals that a purely lysogenic strategy is favored given low cell abundances and high resources (above the (*S*_*c*_, *R*_*c*_) curve) and a purely lytic strategy is favored given high cell abundances and low resources (below the (*S*_*c*_, *R*_*c*_) curve). Analogous to the resource-implicit case, these bounds provide the basis for identifying feasible strategies, depending on whether prophage provide a direct benefit or impose a cost to cellular fitness.

First, consider the case where prophage provide a direct benefit to cellular fitness, *i.e.*, such that (*b*_*L*_/*d*_*L*_) *>* (*b*_*S*_/*d*_*S*_) at the virus-free equilibrium. It would seem apparent that lysogeny (and therefore temperate strategies) should enable viral invasion of an entirely susceptible host population. Indeed, a purely vertical strategy is feasible irrespective of cell density because newly produced lysogens out-compete resident cells; this is true for both resource-implicit (see Figures 3A and 3C) and resource explicit models (see Figure 4A). In contrast, the 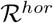 of a horizontal strategy increases with increasing susceptible density *S*^∗^, such that it becomes feasible at a critical value *S*^∗^ = *S*_*c*_ for resource-implicit models (see Figure 3) or beyond a curve (*R*_*c*_, *S*_*c*_) for resource-explicit models (see Figure 4). As a consequence, intermediate temperate strategies with 0 *< p <* 1 can also be feasible at both low and high extremes. In this case, temperate viruses derive most of their fitness via vertical transmission when *S*^∗^ is low and via horizontal transmission when *S*^∗^ is high. In practice, we find that temperate strategies can be feasible across the entire range of host densities (and resource levels), including in circumstances where lytic strategies have 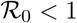 (see Figures 3 and 4).

**FIG. 3:**
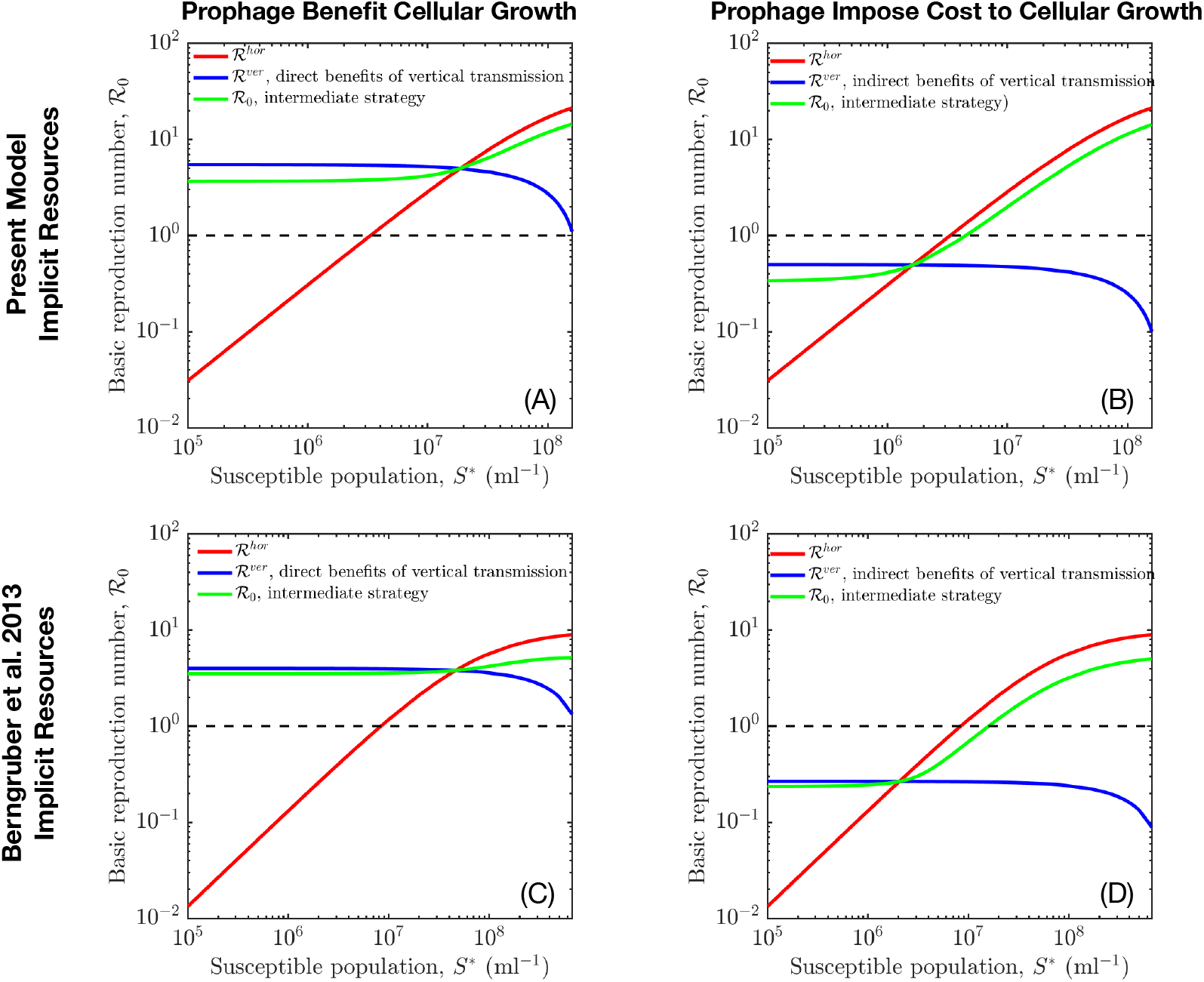
Feasible invasion for viral strategies given variation in susceptible host densities. In panels A and C, prophage provide direct benefit to cellular fitness, *i.e.*, 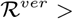 1 given variation in susceptible host densities. In contrast, panels B and D show the case that prophage impose cost to cellular fitness, *i.e.*, 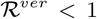 given variation in susceptible host densities. For the intermediate strategy, the probability of lysogeny is *p* = 0.5 and the induction rate is *γ* = 0.1/hr, see model details and relevant parameters in Results, Methods, Appendix A and Appendix E.

**FIG. 4:**
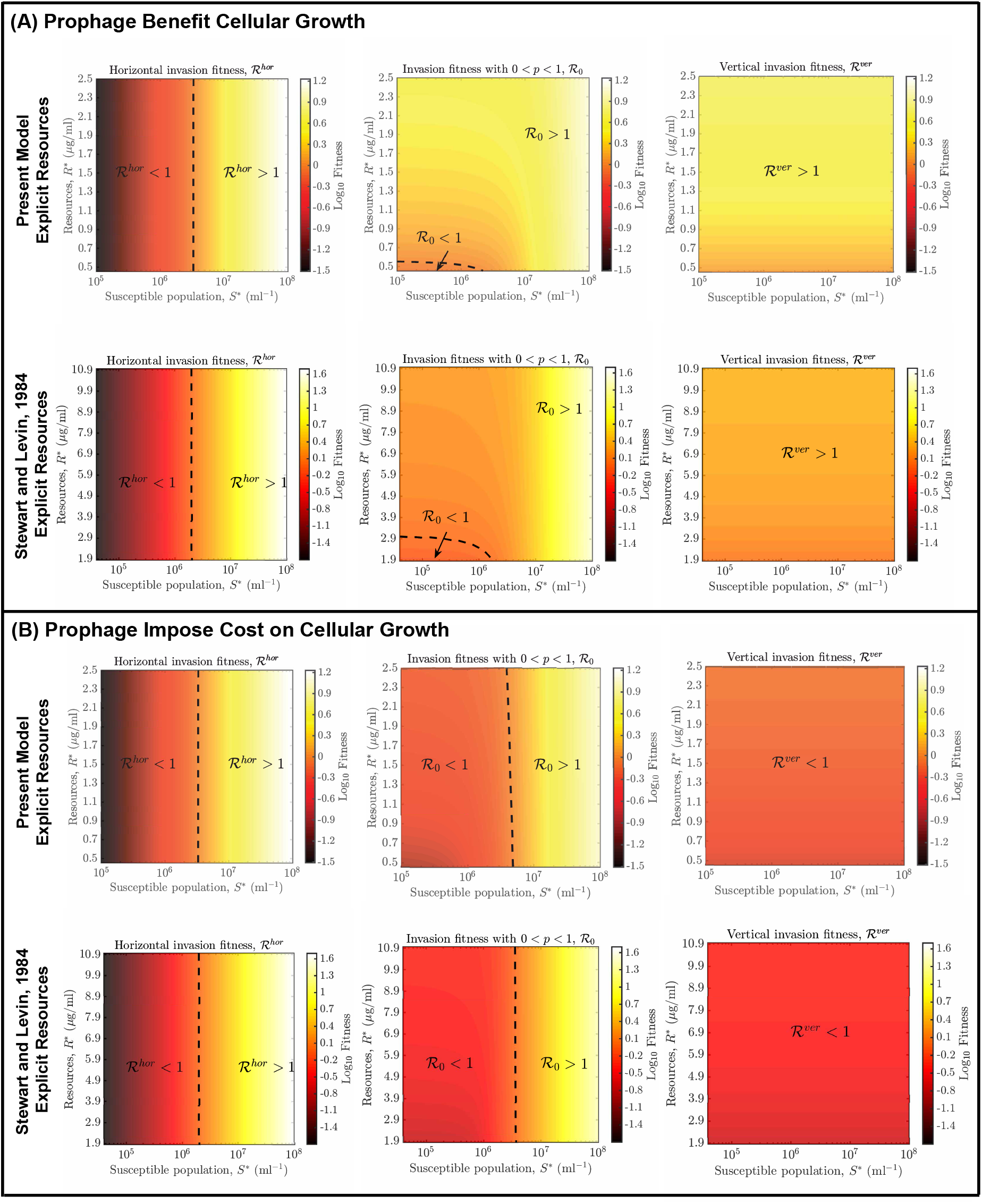
Feasible invasion for viral strategies given variation in resources and susceptible host densities. (A) Prophage provide direct benefit to cellular fitness, *i.e.*, 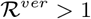 given variation in resources and susceptible host densities. (B) Prophage impose cost to cellular fitness, *i.e.*, 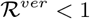 given variation in in resources and susceptible host densities. The probability of lysogeny is *p* = 0.5 and the induction rate is *γ* = 0.1/hr for the intermediate strategy. Additional model details and relevant parameters are in Methods, Appendix A and Appendix E.

In contrast, using the same analysis framework we find that a purely lysogenic strategy with *p* = 1 is not feasible when prophage impose a cost to cellular fitness, *i.e.*, (*b*_*L*_/*d*_*L*_) *<* (*b*_*S*_/*d*_*S*_), at the virus-free equilibrium. This holds both for the resource-implicit and resource-explicit models (see Figures 3B, 3D and 4B). As a result, a temperate strategy with *p >* 0 is feasible insofar as there are sufficiently high host densities for viruses to spread predominantly via a horizontal route. In that case, a sufficient condition for temperate strategy invasion is that 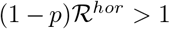, such that the range of ecological conditions for feasible temperate strategies is more restricted than for purely lytic strategies (see Figures 3 and 4).

In summary, temperate phage that confer a benefit to cellular fitness may invade across a wide range of ecological contexts. Yet the more interesting question raised by this analysis is: can temperate viruses that impose a direct fitness cost to cells nonetheless invade in a greater range of ecological contexts albeit when viruses are present?

### E. Endemically infected states and viral invasion

We examine the extent to which temperate phage can invade environments with pre-existing (*i.e.*, circulating) lytic viruses. We are interested specifically in the case where prophage impose a direct cost to cellular fitness, as measured in terms of the ratio of cellular reproduction to mortality in an otherwise virus-free environment. To do so, we first apply an invasion analysis (22) to the resource-explicit model with explicit infections. We assume the resident strain is a purely viral strategy (*p*_*r*_ = 0, *γ*_*r*_ = *γ*_*min*_). We then determine how a strategy (*p*_*m*_, *γ*_*m*_) of a mutant type competes in the environment determined by the resident. We note that the resident endemic equilibrium is stable given the particular parameter set in Appendix E.

Consider the case when the purely lytic strategy has invaded a region with initially high susceptible host densities (high enough that a purely latent strategy would not feasibly invade – see Figure 5A, first blue diamond). Paradoxically in this case, the environment set by the resident lytic strategy can (but not always) be invaded by a temperate strategy (*i.e.*, 0 *< p*_*m*_ ≤ 1) even if the temperate phage imposes a direct cellular fitness cost. The reason is as follows. Initially in the virus-free environment, the susceptible host density is high and resources are low. Lysis depletes susceptible hosts, which reduces niche competition between cells, thereby increasing the potential benefits of vertical transmission. This feedback implies that the vertical fitness can be higher than the horizontal fitness in the environment that lytic phage established, enabling more temperate strategies to invade (see Appendix D for mathematical details). We also note that invasion by a mutant strain does not necessarily imply replacement of the resident strain. If the lytic virus were to become rare, then the lysogens would be out-competed by uninfected hosts, transforming the environment into one susceptible to proliferation by lytic viruses. As such, temperate viruses can invade and then coexist with lytic viruses (see Figure 5A).

**FIG. 5:**
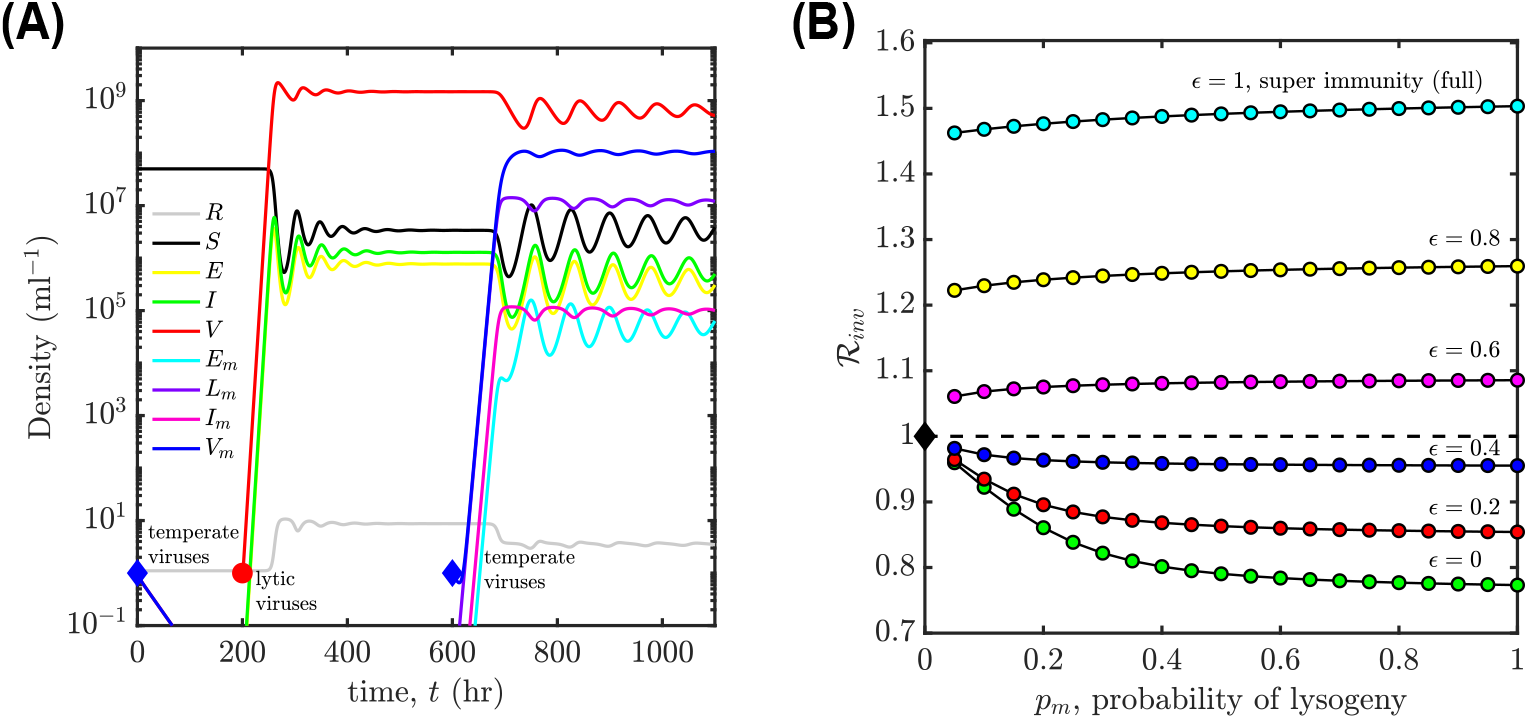
Invasion of temperate phage with varying degrees of super-infection immunity from *ϵ* = 0 (none) to *ϵ* = 1 (complete). (A) Example time series of population densities in the case that lytic phage can drive down microbial cell densities so as to enable invasion by temperate phage. Concretely, at time *t* = 0 hr, temperate phage with purely lysogenic strategy cannot invade virus-free environment (first blue diamond); at time *t* = 200 hrs, purely lytic phage invade virus-free environment (red circle) and spread; at time *t* = 600 hrs, the endemic environment set by the resident lytic strategy can be invaded by temperate phage with purely lysogenic strategy with full immunity *ϵ* = 1 (see second blue diamond). (B) The invasion fitness 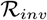 of a temperate phage strategy *p*_*m*_ given variation in *ϵ* from *ϵ* = 0 to *ϵ* = 1, the induction rate is fixed, *γ*_*m*_ = 10*−*2/hr. The model details are presented in Eqs. 1 and Appendix D3, the relevant parameters are selection coefficient *α*_*s*_ = 0.42 and decay rate of lysogens *d*_*L*_ = 0.5/hr (in the case when prophage impose cost to cellular fitness), influx rate of resources *J* = 5.33 (*µg*/ml) h^*−*1^, see additional parameters in Appendix E.

Beyond the density-dependent effect, temperate strategies that impose a fitness cost may come with another form of benefit: conferring ‘super-infection immunity’ to infection by lytic viruses. Super-infection denotes the possibility that a cell is infected by more than one virus (23). Super-infection immunity becomes relevant from an ecological perspective when the resident virus reaches relatively high densities, *i.e.*, in the endemic context. Hence, in an effort to generalize the dynamical example in Figure 5A, we consider a spectrum of cases in which lytic viruses are absorbed into all cells but only infect a fraction (1 − *ϵ*) of lysogens, where *ϵ* is interpreted as the degree of super-infection immunity (see Appendix D3 for details). Hence, a temperate viral strategy is represented by the combination: (*p, γ, ϵ*). We then measure the invasion fitness 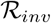 of a temperate virus strategy given variation in *ϵ* from *ϵ* = 0 (all lysogens infected by lytic viruses, no superinfection immunity) to *ϵ* = 1 (no lysogens infected by lytic viruses, full superinfection immunity). We find that temperate phage can invade an environment with hosts and a resident lytic virus insofar as they provide a critical level of super-infection immunity (see Figure 5B).

## III. DISCUSSION

We have demonstrated the benefits of being temperate by exploring the dependency of viral invasion fitness on infection mode and ecological context. By integrating a cell-centric metric of viral invasion fitness (16) and Levins’ loop analysis (17), we derived an interpretable representation of invasion fitness 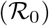 common to a series of mechanistic representations of temperate phage (see boxed Eq. [13] and Figure 2). In contrast to the previous application (21) of loop analysis, our work shows that the loop interpretation of 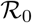 is useful even though the NGM has multiple non-zero eigenvalues. Using this form, we have shown that temperate strategies are feasible (*i.e.*, 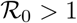) for a wide range of ecological conditions, insofar as prophage provide a direct benefit to cellular fitness. In contrast, we find that temperate phage that impose a direct cost to cellular fitness can still invade environments with pre-circulating lytic viruses (particularly when they provide a critical level of superinfection immunity).

The Levins’ loop approach also enables the interpretation of temperate phage fitness, at least in the short-term. In the short term, the invasion fitness 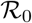 depends on the infection mode as well as an ecological context, e.g., susceptible cell density and resources. The invasion fitness reflects contributions from vertical, horizontal, and mixed transmission pathways. The purely vertical fitness increases as resources increase or host abundances decrease, see Figure 3. In contrast, the purely horizontal fitness increases as host abundances increase, see Figure 3. As a consequence, we find that temperate phage feasibility is enhanced given low host abundances and high resources, whereas the purely lytic strategy is infeasible for low host abundances. This relationship is consistent with observations that lysogeny is prevalent at times of low resources and low host availability, e.g., in aquatic systems (24, 25, 26). In practice, nutrient concentrations co-vary with host cell abundances, and so disentangling the effects of fluxes vs. pools will be critical to translating findings here to analysis of latency in the environment (27, 28, 29, 30, 31).

As a step toward understanding long-term evolution of temperate strategies, we considered the invasibility of endemic states, *i.e.*, by examining whether temperate phage can invade an ecological context with a circulating lytic phage. We find that such endemic virus environments are often invasible by temperate strategies (*i.e.*, 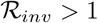) even when prophage impose a cost to cellular growth. This invasibility arises because lytic infections decrease susceptible host cell densities, and indirectly reduce niche competition between uninfected cells and a subpopulation of lysogens. Hence, our results provide direct support for the low host cell density adaptation hypothesis. However, our results also go further. We also evaluated the sensitivity of model findings to variation in superinfection exclusion. We find that when temperate phage impose a cost to cellular growth but confer immunity to lysogens against infection by lytic viruses, then temperate phage can invade in ecological contexts with pre-existing viruses where they would otherwise not be able to invade in a virus-free case. Hence, lytic viruses may actually enable the invasion of a broader range of temperate strategies. This finding provides additional mechanistic support consistent with studies of the evolution of viral strategies given long-term viral-host feedback (15, 32, 33). Note that such invasions are a first-step towards understanding long-term evolution. Indeed, subsequent invasions of an endemically infected state by temperate or purely lytic phage could lead to multi-strain coexistence – further complicating the benefits of being temperate.

While our study identifies possible conditions under which latency may evolve in phage-bacteria communities, additional theory is needed that explicitly models the long-term evolution of transmission strategies in viruses. For example, our analysis assumes that resources affect cell growth but not viral fitness. However, viral life history traits can, in fact, depend on resource levels and host physiological state (34, 35). Hence, future research should connect these microscopic traits with population level models. In addition, infected cell fate is strongly influenced by the cellular multiplicity of infection (cMOI), *i.e.*, the number of co-infecting phage genomes in an individual cell. For phage *λ*, the fraction of lysogeny increases with increasing cMOI (36, 37, 38). The mechanistic basis for this change has focused on feedback in the cell fate determination circuit (39, 40, 41). However, the eco-evolutionary question of why such features have evolved remains an open question. Moving forward, the integration of resource dynamics, multiple infections, and fluctuating ecological dynamics (42) are critical to a comprehensive understanding of why phage are temperate. In doing so, it will also be important to recognize that what may be evolutionary ‘optimal’ in a single-host environment will likely differ in a complex community when multiple hosts and viruses interact (4). For example, a lysogen may not necessarily confer immunity to co-circulating viral types, providing potential advantages to lytic phage in complex communities relative to those in single host-virus populations.

In closing, the loop-based analysis developed here provides a unified and tractable interpretation of temperate phage invasion fitness. Lysogeny provides a direct fitness benefit to viruses when hosts are rare (but resources are available) and also enables viruses to invade environments in which lytic viruses have reduced host densities and by extension niche competition. We speculate that a loop-based approach to measuring invasion fitness may be of service in the analysis of other parasite-host systems. Altogether, our results provide a principled framework for connecting intracellular exploitation of bacteria by phage with population and evolutionary-level outcomes. We hope this framework can facilitate analysis of the experimental evolution of latency in model phage-bacteria systems as well as shed light on drivers of variation in lysogeny in the environment.

## IV. METHODS

### A. Main models

We represent the models in Figure 1 in terms of systems of nonlinear ordinary differential equations (ODEs). Note that the nonlinear population models with explicit infections (both resource-implicit and resource-explicit) are presented in Results. The resource-implicit model with implicit infections given by Berngruber et al., 2013 (15) is

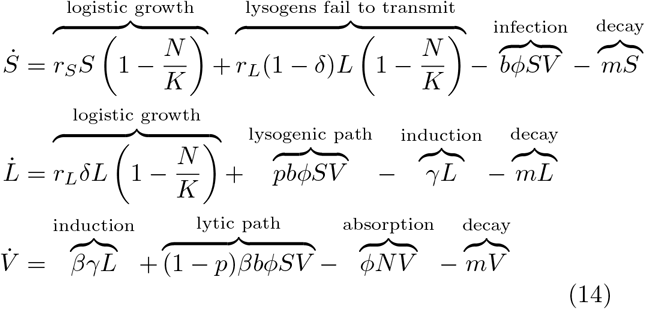

where *N* = *S* + *L* is the density of total cells, *L* is the density of infected cells, the susceptible cells density is *S* and the free-virus density is *V*. More details can be found in (15).

The resource-explicit model with implicit infections given by Stewart & Levin, 1984 (13) is

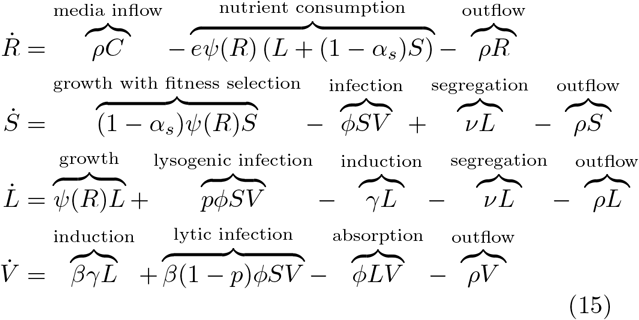

where *R*, *S*, *L*, and *V* denote the densities of resources, susceptible cells, lysogens and virus particles, respectively. This model describes the dynamics of microbial populations in the chemostat, *ρ* is the inflow (and outflow) rate; additional model details in (13).

### B. Viral invasion analysis

The 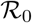 calculations for each model variant closely follow the procedures given by Diekmann (18); complete calculations are provided in Appendix B. To demonstrate the biological interpretation of NGM associated with temperate phage invasion dynamics, we present the NGM, Φ, of system of Eqs. 1 in Eq. [5]. The loop-based 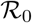 interpretation is inspired by Levins’ loop analysis (17). For each model, we first compute 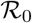 via Eq. [9] by an NGM approach, then, reformulate it into Eq. [13] by identifying all the one- and two-generation loops. The loop-based results are summarized in Table (I) and the detailed calculations are given in Appendix B.

### C. Endemic invasion analysis

For each temperate phage model, we construct the mutant-resident system, and compute the invasion fitness of mutant viral strains 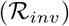 at the resident endemic equilibrium via a next-generation matrix approach (22, 32); see Appendix D for details.

### D. Data availability

Simulation code is written in MATLAB and available at https://github.com/WeitzGroup/Guanlin_when_be_temperate.

## Acknowledgments

We thank An Qi for reviewing code. This work was supported by a grant from the Simons Foundation (SCOPE Award ID 329108, J.S.W.).

## Supplementary Information

### A Nonlinear, Population Model of Temperate Phage Dynamics

#### A.1 Main models

The resource-implicit model with explicit infections from the main text is

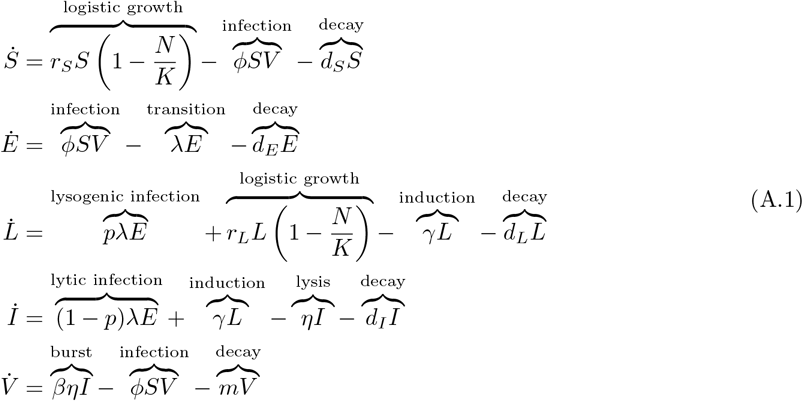

where *S*, *E*, *L*, *I* and *V* denote the densities of susceptible cells, exposed infected cells, lysogens, lytic-fated infected cells and virus particles respectively, and *N* = *S* + *E* + *L* + *I* is the total cell density. Parameters *r*_*S*_ and *r*_*L*_ denote the maximal cellular growth rates of susceptible cells and lysogens, *K* is the carrying capacity, *φ* is the adsorption rate, *d*_*S*_, *d*_*E*_, *d*_*L*_ and *d*_*I*_ are the cellular death rates of susceptible cells, exposed infected cells, lysogens and lytic-fated infected cells respectively, *λ* is the transition rate from exposed cells to the fate determined cells, *p* is the probability of lysogenization, *γ* is the induction rate, *η* is the lysis rate, *β* is the burst size and *m* is the virion decay rate.

The resource-explicit model with explicit infections from the main text is

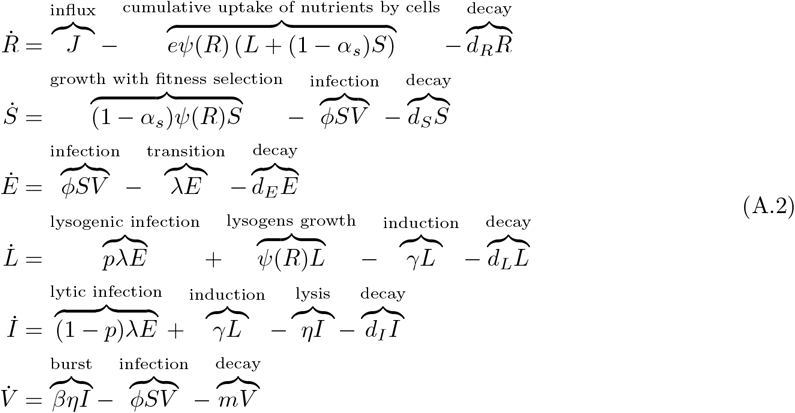

where *R*, *S*, *E*, *L*, *I* and *V* denote the densities of resources, susceptible cells, exposed infected cells, lysogens, lytic-fated infected cells and virus particles, respectively. The growth function *ψ*(*R*) = *µ*_*max*_*R*/(*R*_*in*_ + *R*) is a Monod equation, where *µ*_*max*_ is the maximal cellular growth rate and *R*_*in*_ is the half-saturation constant. The parameters *J* and *d*_*R*_ are the influx and decay rates of resources, *e* is the host conversion efficiency, *α*_*S*_ is the selection coefficient that measures the relative fitness between lysogens and susceptible cells. All other parameters are defined as in model [A.1]. The resource-implicit model with implicit infections in Berngruber et al. (2013) is

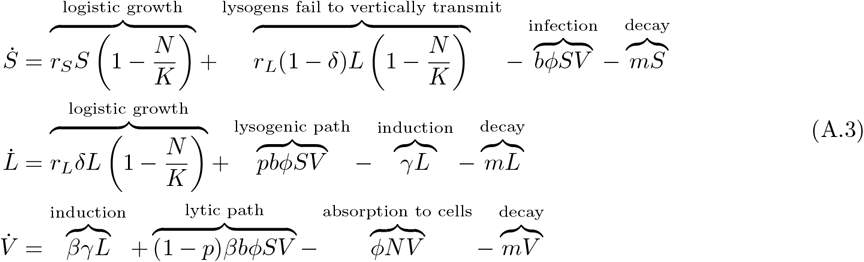

where *N* = *S* + *L* is the density of total cells, *L* is the density of infected cells, the density of susceptible cells is *S* and the free-virus density is *V*. More details about model [A.3] can be found in ([1]).

The resource-explicit model with implicit infections from Stewart and Levin (1984) is

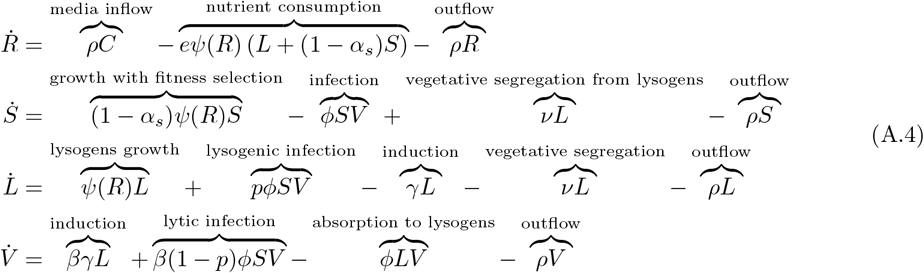

where *R*, *S*, *L*, and *V* denote the densities of resources, susceptible cells, lysogens and virus particles, respectively. Model [A.4] describes the dynamics of populations in a chemostat, where *ρ* is the inflow (and outflow) rate and *ν* is the segregation rate whereby lysogens become susceptible cells. See ([2]) for more details.

In the four model variants, the phage strategies are defined by two parameters: *p* defines the probability a virus enters the lysogenic pathway and *γ* defines the induction rate after a virus enters the lysogenic pathway. We define the viral strategy space Θ as

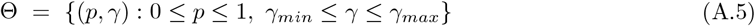

where *γ*_*min*_ > 0.

#### A.2 From explicit infections to implicit infections

In this section, we show that the explicit infection models [A.1] and [A.2] can be reduced to the models with implicit infections via a quasi–steady–state (QSS) approximation.

For model [A.1] and model [A.2], we assume the (lysis-lysogeny decision) transition process and lysis process are extremely rapid in comparison to all the other biological processes. In other words, we let *λ* ≫ *d*_*E*_ and *η* ≪ *d*_*I*_. The population dynamics of exposed cells from model [A.1] and model [A.2] can be rewritten as

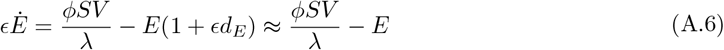

where *ϵ* = 1/*λ* ≪ 1. Hence, the QSS equilibrium density of exposed cells population is *E*^*ϵ*^ = *φSV/λ*. Using *E*^*ϵ*^, the population dynamics of lytic-fated infected cells from model [A.1] and model [A.2] are rewritten as

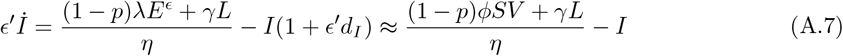

where *ϵ′* = 1/*η* ≪ 1. Hence, the QSS equilibrium density of lytic-fated infected cells is *I*^*ϵ′*^ = [(1 − *p*)*φSV* + *γL*]/*η*. Substituting *E*^*ϵ*^ and *I*^*ϵ′*^ into model [A.1] reduces it to the *S, L, V* -system with implicit infections,

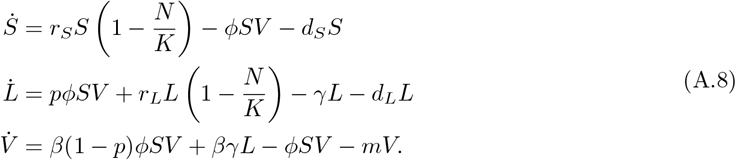

Substituting *E*^*ϵ*^ and *I*^*ϵ′*^ into model [A.2] reduces it to the *R, S, L, V* -system with implicit infections,

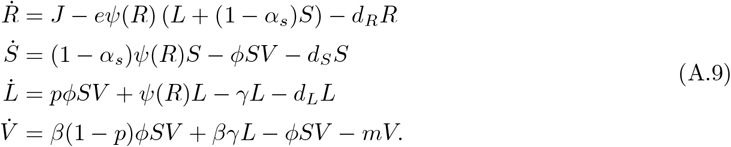

### B Viral Invasion Analysis

#### B.1 Virus-free equilibrium

For the resource-implicit models [A.1] and [A.3], there are only susceptible cells (*S*^∗^) in the virus-free environments. The virus-free equilibrium of model [A.1] is

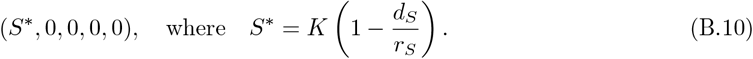

The virus-free equilibrium of model [A.3] is

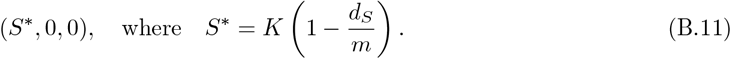

For the resource-explicit models [A.2] and [A.4], there are both resources and susceptible cells (*R*^∗^, *S*^∗^) in the virus-free environments. The virus-free equilibrium of model [A.2] is

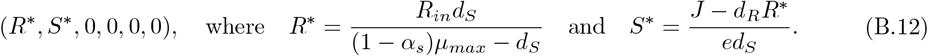

The virus-free equilibrium of model [A.4] is

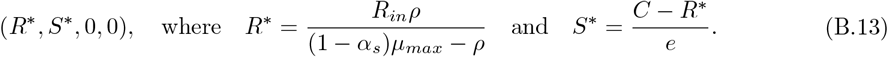

#### B.2 Next-generation matrix approach

We start by computing the next-generation matrix (NGM) for the resource implicit model with explicit infections [A.1]. Consider the Jacobian of the model [A.1] evaluated at the virus-free equilibrium, Eq. [B.12]. We denote *𝒥* as the submatrix of Jacobian for the *E, L, I, V* -subsystem. We decompose the submatrix as *𝒥* = *F* + *𝒱* where the transmission matrix *F* and the transition matrix *𝒱* are

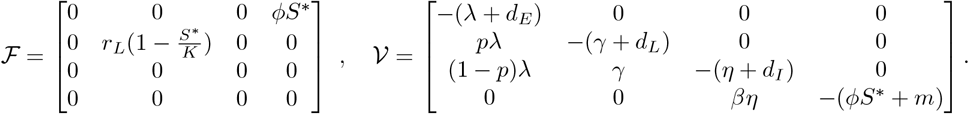

Via NGM theory ([3]), the basic reproduction number 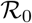 corresponds to the largest eigenvalue of the matrix 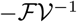, namely 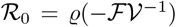 where *ϱ*(*M*) is the spectral radius of the matrix *M*. There are two epidemiological birth states in model [A.1], hence, there are only two non-zero eigenvalues in 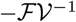. Here we introduce an augmented operator *Q*, where *Q* is a matrix in ℝ^4*×*2^ with unit vectors in columns 1 and 2. The spectral radius of the 4 *×* 4 matrix 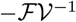 is the same as the spectral radius of the 2 *×* 2 matrix 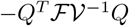. For convenience, we define the next generation matrix of model [A.1] as

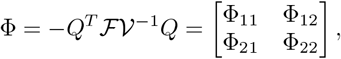

where the entries of Φ are

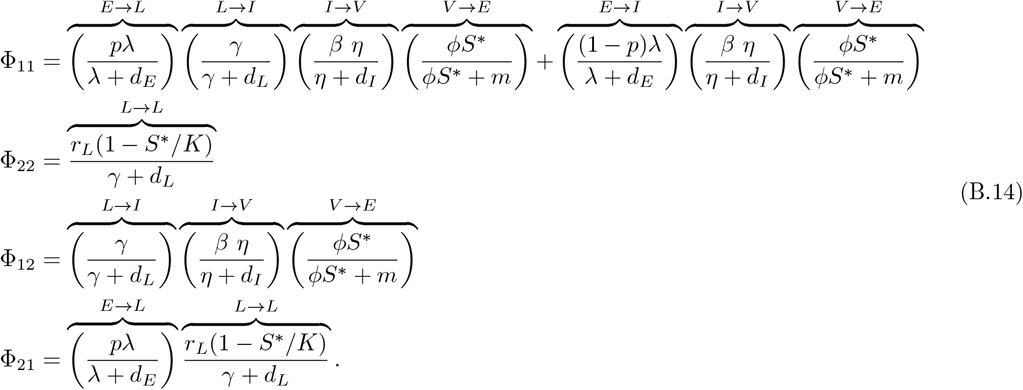

Entry Φ_*ij*_ represents the expected number of new infected individuals in state *i*, generated by one infected individual at state *j* (*i, j* = *L, E*), accounting for new infections that arise via the lytic and lysogenic pathways. Φ can be rewritten in terms of the basic reproductive number for viruses with purely lytic strategies (*p* = 0; 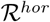) and the basic reproductive number for viruses with purely lysogenic strategies (*p* = 1, *γ* = 0; 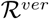),

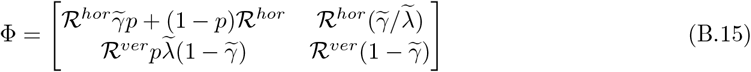

where

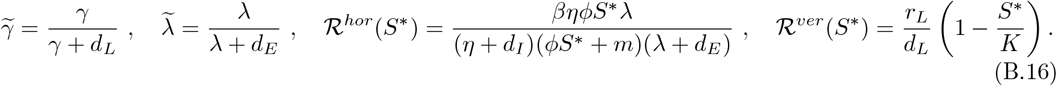

Next we compute the NGM for the implicit model with implicit infections [A.3]. We decompose the linearized infected subsystem of model [A.3], at the virus-free equilibrium, Eq. [B.11], into the transmission matrix 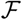 and the transition matrix *𝒱*:

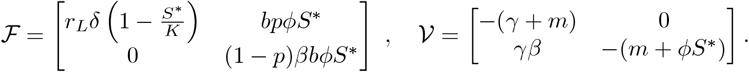

The NGM of model [A.3] is defined as

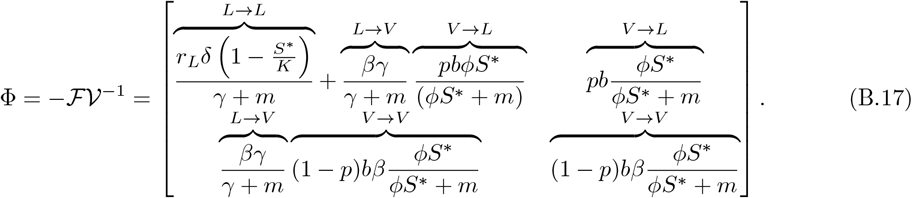

As before, the NGM can be rewritten as

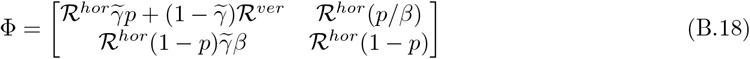

where

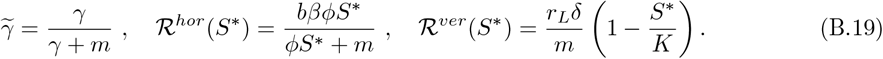

Similarly, we compute the NGM for the resource explicit model with explicit infections [A.2]. We decompose the linearized infected subsystem of model [A.2] evaluated at the virus-free equilibrium, Eq. [B.12], into the transmission matrix 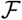 and transition matrix *𝒱*:

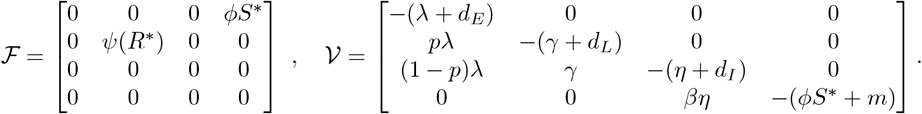

The NGM of model [A.2] is defined as

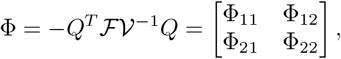

where *Q* is a matrix in ℝ^4*×*2^ with unit vectors in columns 1 and 2. The entries of Φ are

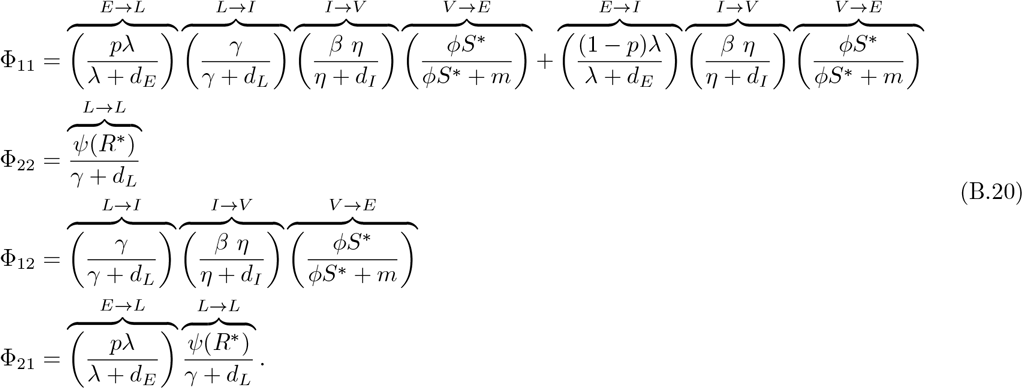

As before, we can rewrite the NGM as

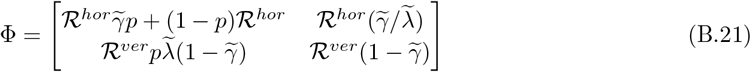

where

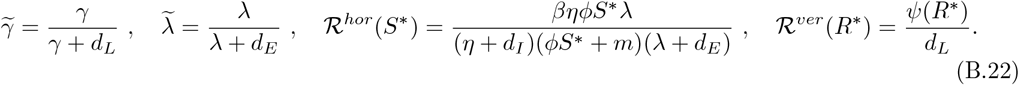

Finally, we compute the NGM for the resource explicit model with implicit infections [A.4]. We decompose the linearized infected subsystem of model [A.4], at the virus-free equilibrium, Eq. [B.13], in to the transmission matrix 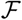 and the transition matrix *𝒱*:

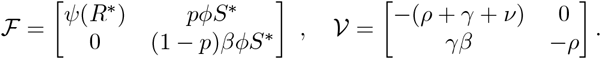

The NGM of model [A.4] is defined as

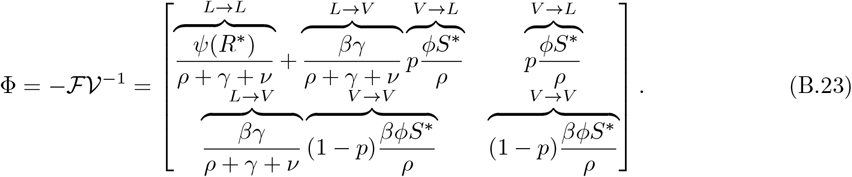

As before, we can rewrite the NGM as

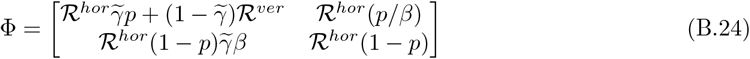

where

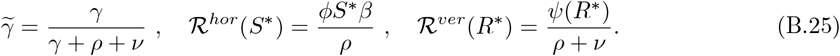

Notably, the next-generation matrices for all four model variants are 2 × 2 matrices, as shown in Eq. [B.15], Eq. [B.17], Eq. [B.21] and Eq. [B.24]. The traces, *T* (Φ), and determinants, *D*(Φ), of each next-generation matrix can be written as

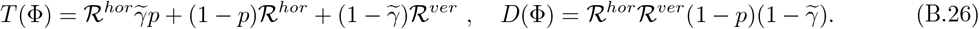

where the values of 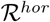, 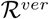 and 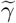 are given in Eq. [B.16], Eq. [B.19], Eq. [B.22] and Eq. [B.25]. The dominate eigenvalue 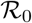 can be obtained from the trace and the determinant of the next-generation matrix,

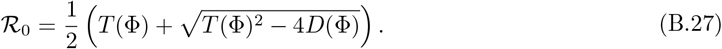

In general, 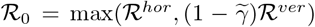 when *p* = 0. However, we define 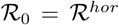 when *p* = 0 because that matches the biological scenario defined by a virus with a purely lytic strategy.

#### B.3 Loop-based interpretation of 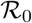

Motivated by Levins’ loop analysis ([4]), we can rewrite Eq. [B.27] in a biological interpretable way. We define, 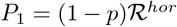, 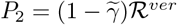 and 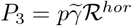. Then, Eq. [B.27] can be written as

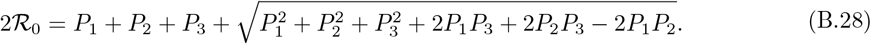

A full description of this interpretation is presented in the main text.

### C Maximization of 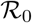 and Feasible Invasion Strategies

#### C.1 Preliminaries

A strategy (*p*^∗^, *γ*^∗^) maximizes the basic reproduction number corresponding to the viral invasion fitness as measured at the individual level if

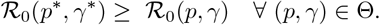

Given its use in the calculations and proofs, we note that while 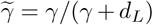 lies in the closed interval

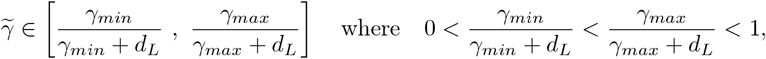

we generally use 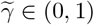 instead of *γ ∈* [*γ*_*min*_, *γ*_*max*_] for convenience.

**Proposition C.1** *The basic reproduction number* 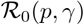, *Eq. [B.27], is imarginal continuous on* (*p, γ*) ∈ *Θ and marginal differentiable on* (*p, γ*) *∈* Θ *except at* 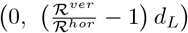.

**Proof.** The marginal continuity of *R*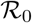(*p, γ*) on (*p, γ*) ∈ Θ is obvious. Let 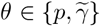, the derivative of 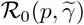 with respect to *θ* is

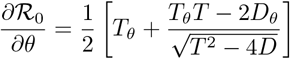

where 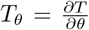 and 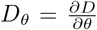. Thus, we need *T*^2^ *−* 4*D* to be strictly positive to ensure the marginal differentiability. We write out *T*^2^ *−* 4*D* in terms of *p* and 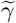

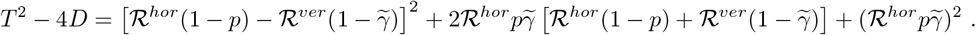

Algebraic manipulation shows that *T*^2^ *−* 4*D >* 0 everywhere in Θ except at 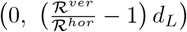. □

##### Lemma C.2

*Given a fixed γ satisfying* 0 < *γ*_*min*_ ≤ *γ* ≤ *γ*_*max*_, *the basic reproduction number* 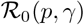 *is uniquely maximized at p*^∗^ = 1 *if* 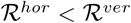. *In contrast, the basic reproduction number* 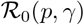 *is uniquely maximized at p*^∗^ = 0 *if* 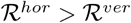.

**Proof.** In the proof, we assume 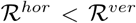; the case 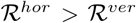 can be proved with a similar argument. The uniqueness of the maximal strategy *p*^∗^ is showed by the strict monotonicity of 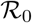 for all *p* ∈ [0, 1]. Calculating the derivative of 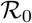 with respect to *p* yields

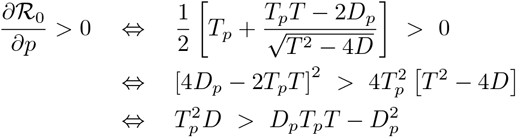

where the derivatives are 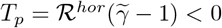 and 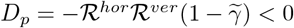. Note that

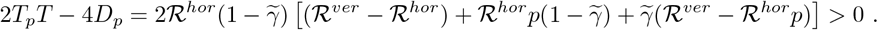

Substitution yields

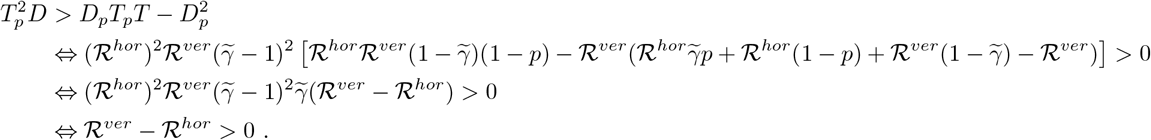

Therefore, the basic reproduction number is a strictly monotone increasing function of *p* in the range of [0, 1]. This implies 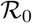 is maximized at *p*^∗^ = 1 and minimized at *p* = 0 by the continuity of 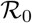. □

##### Lemma C.3

*Given a fixed p satisfying* 0 *< p ≤* 1, *the basic reproduction number* 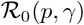 *is uniquely maximized at* 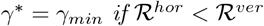. *In contrast, the basic reproduction number* 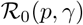 *is uniquely maximized at 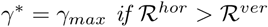*.

**Proof.** The proof is similar to the proof of lemma (C.2). According to proposition (C.1), we fix *p* ≠ 0 such that 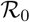 is differentiable on *γ*_*min*_ ≤ γ ≤ *γ*_*max*_. We assume 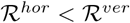; the proof for 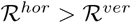 is similar. Note by the chain rule that

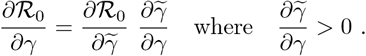

Let 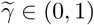. Differentiating 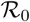 yields

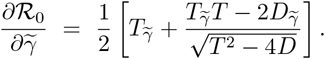

Given 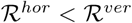, the derivatives are 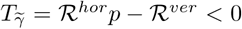 and 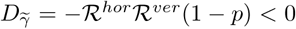. The sign of 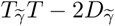 is unknown, which can be expanded as

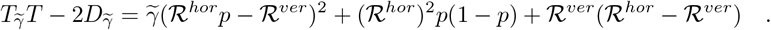

If 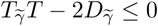 then 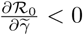. If 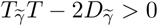, then

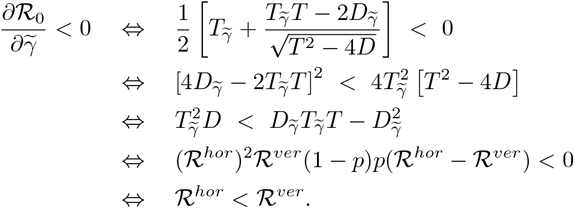

Thus, if 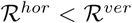 then 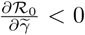. This means the basic reproduction number is a strictly monotone decreasing function of *γ* when 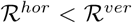. Thus, by the continuity of 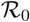 is maximized at *γ*^∗^ = *γ*_*min*_ and minimized at *γ* = *γ*_*max*_ uniquely. □

#### C.2 Maximization of 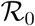 and feasible invasion strategies

Using lemma (C.2) and lemma (C.3), we summarize the results of maximal viral strategy in the following theory.

##### Theorem 1

*The basic reproduction number* 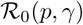(*p, γ*), *Eq. [B.27], is maximized at p*^∗^ = 0 *and all γ*_*min*_ ≤ *γ* ≤ *γ*_*max*_ *if* 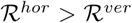. *On the other hand*, 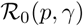 *is maximized uniquely at* (*p*^∗^, *γ*^∗^) = (1, *γ*_*min*_) *if* 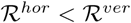.

**Proof.** The proof follows from lemma (C.2) and lemma (C.3). If 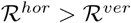, then

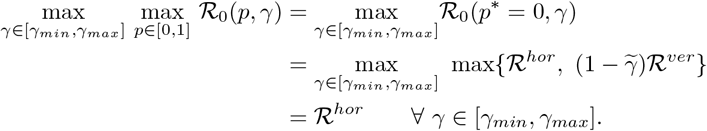

On the other hand, if 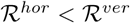, then

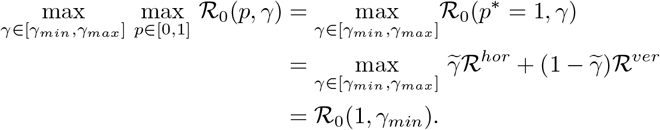

□

**Remark 1** *We note the following*:

*(i)* 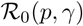 *is bounded above by* 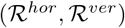 *and bounded below by* 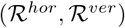.

*(ii) If* 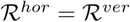, *then* 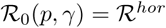 *for all* (*p, γ*) ∈Θ.

*(iii) If* 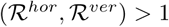, *then* 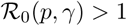 *for all* (*p, γ*) ∈Θ.

*(iv) If* 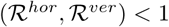, then 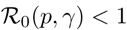 *for all* (*p, γ*) ∈Θ.

*(v) If* 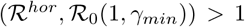 *and* 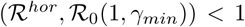, *then there exists a critical curve* 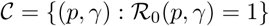, *which partitions the viral strategy space* Θ *into feasible strategy regime (i.e.*, 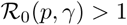*) and infeasible strategy regime (i.e.*, 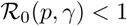).

**Remark 2** *For the two resource-implicit models [A.1] and [A.3], theorem (1) can be restated in terms of the density of susceptible hosts at the virus-free equilibrium (S^∗^). Specifically, because 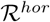 is a monotonically increasing function of S^∗^ and 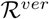 is a monotonically decreasing function of S^∗^, there exists a critical value S_c_ that satisfies 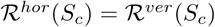. Thus, we can restate theorem (1) in terms of S^∗^: If S^∗^ > S_c_, then 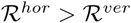 and the maximal strategy is p^∗^ = 0 and arbitrary γ ∈ [γ_min_, γ_max_]. If S^∗^ < S_c_, then 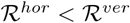 and the maximal strategy is p^∗^ = 1 *and γ* = *γ*_*min*_*.

**Remark 3** *For the two resource-explicit models [A.2] and [A.4], theorem (1) can be restated in terms of the resource concentration and the density of susceptible hosts at the virus-free equilibrium (R^∗^, S^∗^). Specifically, because 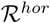 is a monotonically increasing function of S^∗^ and 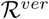 is a monotonically increasing function of R^∗^, there exists a curve in R^∗^, S^∗^-space where there is a transition from the purely lytic strategy (p = 0) maximizing 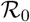 to the purely lysogenic strategy (p = 1, γ = γ_min_) maximizing 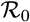. That curve is defined by 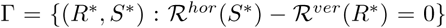. We decompose R^∗^, S^∗^-space into two regions based on Γ, let 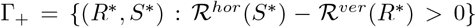 and 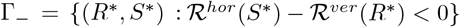. Thus, we can restate theorem (1) in terms of R^∗^ and S^∗^: If (R^∗^, S^∗^) ∈ Γ_+_, then 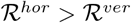 and the maximal strategy is p^∗^ = 1 and arbitrary γ ∈ [γ_min_, γ_max_]. If (R^∗^, S^∗^) ∈ Γ_−_ then 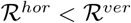 and the maximal strategy is p* = 1 γ = γ_min_*.

**Remark 4** *Let σ denote the largest eigenvalue of the linearized infected subsystem, 𝒥. Biologically, σ is the exponential growth rate of viral strategy in a virus-free host population. Both σ and 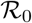 can be used to predict viral invasion because sign 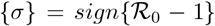. The values of σ and 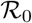 may not be maximized at by the same viral strategy. We numerically explored what viral strategies maximized σ in model [A.1]. Our numerical results showed that there exists a critical density of susceptible hosts, 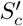, such that if 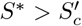 then σ is maximized at p^∗^ = 0 and arbitrary γ ∈ (γ_min_, γ_max_) and if 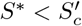 then σ is maximized at p^∗^ = 1 and γ* = *γ_min_. However, the critical values S_c_ and 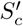 are not the same; see Table (1) for an example. Thus, in general, the viral strategy that maximizes 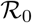 is not the strategy that maximizes instantaneous growth rate*.

### D Endemically Infected States and Viral Invasion

In this section, we provide details about the evolutionary analysis for the four model variants. In doing so, we apply an invasion analyses ([5]), in which the system reaches a stable endemic equilibrium with a resident viral strategy (*p*_*r*_, *γ*_*r*_) ∈ Θ. We then introduce a mutant viral strain with strategy (*p*_*m*_, *γ*_*m*_) ∈ Θ and determine if the mutant can invade the resident equilibrium. We start by deriving the next-generation matrix of the mutant at the resident equilibrium.

#### D.1 Mutual invasion analysis via NGM approach

The mutant-resident system of model [A.1] is

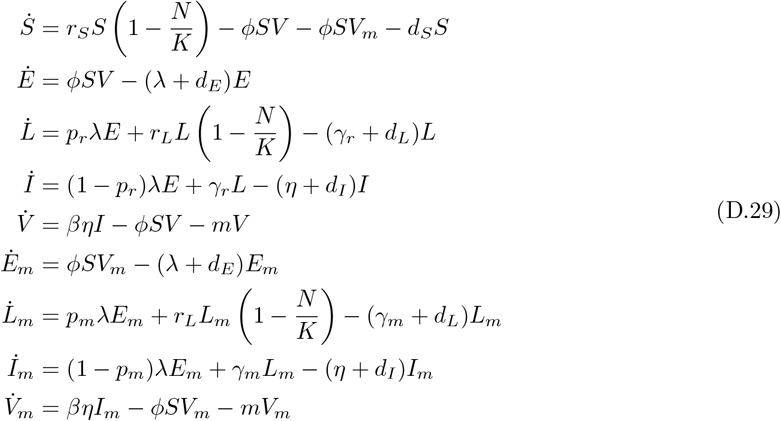

where *N* = *S* +*E* +*L*+*I* +*E*_*m*_ +*L*_*m*_ +*I*_*m*_ is the total population of hosts. The *E*_*m*_, *L*_*m*_, *I*_*m*_, *V*_*m*_-subsystem denotes the classes with the mutant viral strategies. The resident endemic equilibrium of system [D.29] is (*S*^+^, *E*^+^, *L*^+^, *I*^+^, *V*^+^, 0, 0, 0, 0). Invasion of the resident endemic equilibrium is determined by the magnitude of the maximum eigenvalue of the next generation matrix,

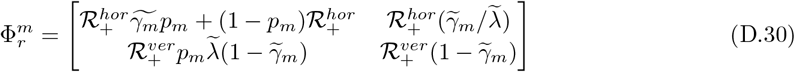

where

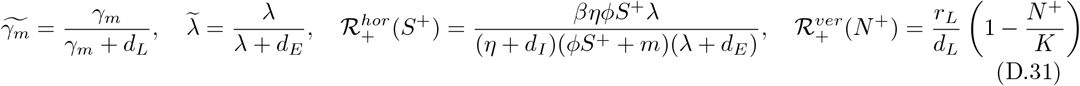

and *N* ^+^ = *S*^+^ + *E*^+^ + *L*^+^ + *I*^+^. The largest eigenvalue of 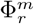 (mutant invasion fitness) is denoted by 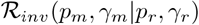. The sign of 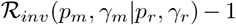 determines the invasion of the resident endemic equilibrium in system [D.29].

Similarly, we add the *L*_*m*_, *V*_*m*_-subsystem to model [A.3]. The resident endemic equilibrium of the augmented resident-mutant system of model [A.3] is denoted by (*S*^+^, *L*^+^, *V*^+^, 0, 0). The next-generation matrix of a mutant strain with viral strategy (*p*_*m*_, *γ*_*m*_) is

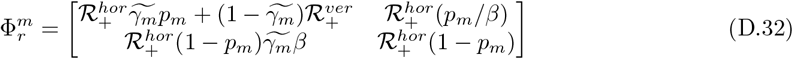

where

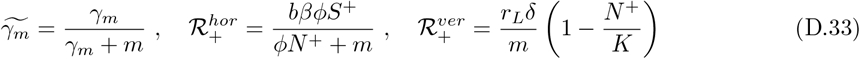

and *N* ^+^ = *S*^+^ + *L*^+^.

Again, we add the *E*_*m*_, *L*_*m*_, *I*_*m*_, *V*_*m*_-subsystem to model [A.2]. The resident endemic equilibrium of the augmented resident-mutant system of model [A.2] is denoted by (*R*^+^, *S*^+^, *E*^+^, *L*^+^, *I*^+^, *V* ^+^, 0, 0, 0, 0). The next-generation matrix of a mutant strain with viral strategy (*p*_*m*_, *γ*_*m*_) is

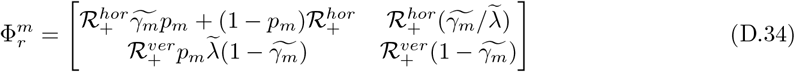

where

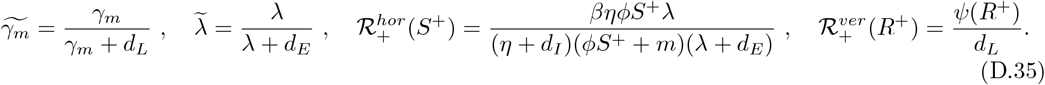

Finally, we add the *L*_*m*_, *V*_*m*_-subsystem to model [A.4]. The resident endemic equilibrium of the augmented resident-mutant system of model [A.4] is denoted by (*R*^+^, *S*^+^, *L*^+^, *V* ^+^, 0, 0). The next-generation matrix of a mutant strain with viral strategy (*p*_*m*_, *γ*_*m*_) is

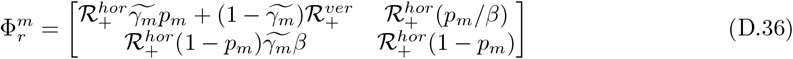

where

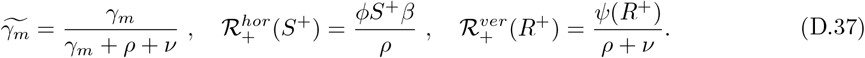

Notably, the derived next-generation matrices for the mutant strains in all four model variants are 2 *×* 2 matrices. The traces, 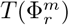, and determinants, 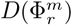 of four next-generation matrices are written as

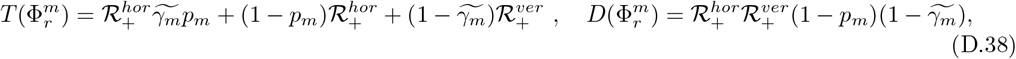

where the 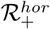, 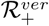 and 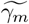 are given in Eq. [D.31], Eq. [D.33], Eq. [D.35] and Eq. [D.37]. The dominate eigenvalue 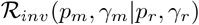 can be obtained from the trace and the determinant of the next-generation matrix,

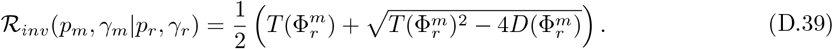

Using the fact that 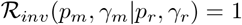 when (*p*_*m*_, *γ*_*m*_) = (*p*_*r*_, *γ*_*r*_), and the monotonicity of 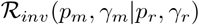 on (*p*_*m*_, *γ*_*m*_) ∈ Θ provided by lemma (C.2), lemma (C.3) and theorem (1), we immediately obtain the following theorem.

##### Theorem 2

*For all of the four model variants, we assume the susceptible cell population is positive at the stable resident endemic equilibrium, i.e., S^+^ > 0. Then, we have*:

i. *If* 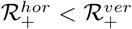, *viral strains with more lysogenic strategies (i.e., p*_*r*_ < *p*_*m*_ ≤ 1 *and γ*_*min*_ ≤ *γ*_*m*_ < *γ*_*r*_) *can invade, and viral strains with more lytic strategies (i.e.*, 0 ≤ *p*_*m*_ < *p*_*r*_ *and γ*_*r*_ < *γ*_*m*_ ≤ *γ*_*max*_) *cannot invade*.
ii. *If* 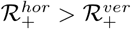, *viral strains with more lysogenic strategies (i.e., p_r_ < p_m_ ≤* 1 *and γ_min_ ≤ γ_m_ < γ_r_) cannot invade, and viral strains with more lytic strategies (i.e.*, 0 ≤ *p_m_ < p_r_ and γ_r_ < γ_m_ ≤ γ_max_) can invade*.

**Remark 5** *For model [A.1] and model [A.2], the stable resident endemic equilibrium might be a boundary equilibrium with S^+^ = 0. If the resident system reaches a stable endemic equilibrium with S^+^ = 0, then no mutant strain can invade. Specifically, a mutant strain is introduced at the resident equilibrium through a virus particle, the first step in the proliferation of a virus is that it infects a susceptible cell, hence, a virus mutant cannot spread given the absence of a susceptible population*.

#### D.2 Robustness of maximal strategies

Theorem (2) requires the specification of the resident equilibrium to determine whether mutant invasion scenarios correspond to case (i) or case (ii). In this section, we provide additional results to apply theorem (2) to model [A.1] and model [A.2]. Further, we assume the resident strain is either the purely lytic (*p* = 0) or purely lysogenic strategy (*p* = 1, *γ* = *γ*_*min*_).

**Proposition D.1** *(i) The stable resident endemic equilibrium of model [A.1]*, (*S*^+^, *E*^+^, *L*^+^, *I*^+^, *V* ^+^) *with S*^+^ *>* 0, *satisfies S*^+^ *< N* ^+^ *< S^∗^, where S^∗^ is given by Eq. [B.10]*.

*(ii) The stable resident endemic equilibrium of model [A.2]*, (*R*^+^, *S*^+^, *E*^+^, *L*^+^, *I*^+^, *V* ^+^) *with S*^+^ *>* 0, *satisfies R*^+^ *> R^∗^ and S*^+^ *< S^∗^, where R^∗^ and S^∗^ are given by Eq. [B.12]*.

**Proof.** Proof of (*i*): *S*^+^ *< N* ^+^ is a trivial observation. We prove *N* ^+^ < *S*^∗^ by contradiction. Suppose *N* ^+^ ≥ *S*^∗^ and set *S*\* = 0 in model [A.1]

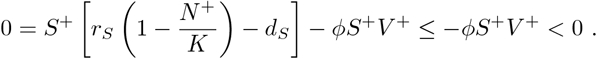

This shows a contradiction such that (*i*) is true.

Proof of (*ii*): At the stable resident endemic equilibrium of model [A.2], we set 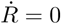 and 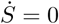

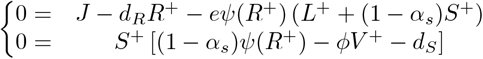

Given *S*^+^ *>* 0, we rewrite the above system and find that

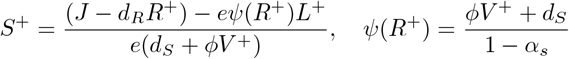

Note that *ψ*(*R*^∗^) = *d*_*S*_ /(1 − *α*_*S*_) by substituting virus-free equilibrium, Eq. [B.12], into the Monod equation. Then, we find that

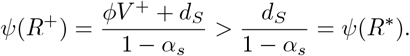

It should be clear that *R*^+^ *> R*^∗^ since *ψ*, the Monod equation, is a strictly monotone increasing function. In addition,

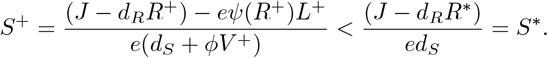

□

**Proposition D.2** *(i) If the virus-free equilibrium of model [A.1] satisfies 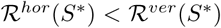, where 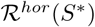 and 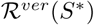 are given by Eq. [B.16], then 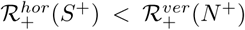 at the stable resident endemic equilibrium, where 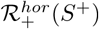 and 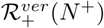 are given by Eq. [D.31]*.

*(ii) If the virus-free equilibrium of model [A.2] satisfies 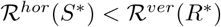, where 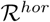 and 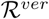 are given by Eq. [B.22], then 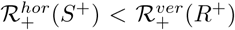 at the stable resident endemic equilibrium, where 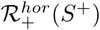 and 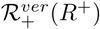 are given by Eq. [D.35]*.

**Proof.** Proof of (*i*): First, we assume that the susceptible cells population at resident equilibrium of model [A.1] is positive, *S*^+^ *>* 0, then we have *S*^+^ *< N* ^+^ < *S*^∗^ by proposition (D.1). Given 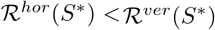 in Eq. [B.16], we arrive

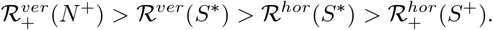

Second, if *S*^+^ = 0 at the resident equilibrium of model [A.1], 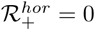, it’s clear that 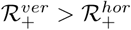.

Proof of (*ii*): First, we assume that the susceptible cells population at resident equilibrium of model [A.2] is positive, *S*^+^ *>* 0, then we have *S*^+^ < *S*^∗^ and *R*^+^ > R^∗^ by proposition (D.1). Given 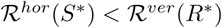 in Eq. [B.22], we arrive

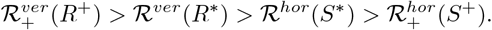

Second, if *S*^+^ = 0 at the resident equilibrium of model [A.2], 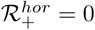, it’s clear that 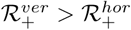. □

For model [A.1] and model [A.2], when the resident strains are either the purely lytic strategy (*p* = 0) or the purely lysogenic strategy (*p* = 1, *γ* = *γ*_*min*_), some qualitative results of evolutionary invasion are immediately obtained by theorem (2) and proposition (D.2). First, in the event that resident strains are purely lytic (*p* = 0), then, we have remark (6).

**Remark 6** *We assume the resident viral strain is the purely lytic strategy (p = 0).*

i. *If the virus-free equilibrium of model [A.1] satisfies 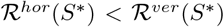, where 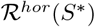 and 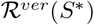 are given by Eq. [B.16], the viral strains with more lysogenic strategies (i.e., 0 < p_m_ ≤ 1) can invade the stable resident endemic equilibrium*.
ii. *If the virus-free equilibrium of model [A.1] satisfies 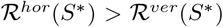, where 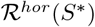 and 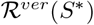 are given by Eq. [B.16], the viral strains with more lysogenic strategies (i.e., 0 < p_m_ ≤ 1) can invade the stable resident endemic equilibrium if 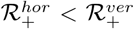, where 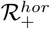 and 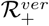 are given by Eq. [D.31]. In contrast, the viral strains with more lysogenic strategies cannot invade the stable resident endemic equilibrium if 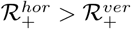*.
iii. *If the virus-free equilibrium of model [A.2] satisfies 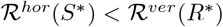, where 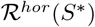 and 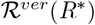 are given by Eq. [B.22], the viral strains with more lysogenic strategies (i.e., 0 < p*_*m*_ *≤ 1) can invade the stable resident endemic equilibrium*.
iv. *If the virus-free equilibrium of model [A.2] satisfies 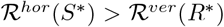, where 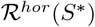 and 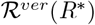 are given by Eq. [B.22], the viral strains with more lysogenic strategies (i.e., 0 < p_m_ ≤ 1) can invade the stable resident endemic equilibrium if 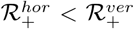, where 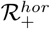 and 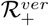 are given by Eq. [D.35]. In contrast, the viral strains with more lysogenic strategies cannot invade the stable resident endemic equilibrium if 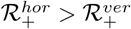*.

Second, in the event that the resident strain is purely lysogenic (*p* = 1, *γ* = *γ*_*min*_) and the susceptible cell population is positive at the resident equilibrium, *S*^+^ *>* 0, then we arrive at remark (7). Note that the invasion scenario of *S*^+^ = 0 is provided in remark (5).

**Remark 7** *We assume the resident viral strain is the purely lysogenic strategy (p = 1, γ = γ_min_) and the susceptible cell population is positive at the resident equilibrium, S^+^ > 0*.

i. *If the virus-free equilibrium of model [A.1] satisfies 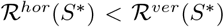, where 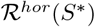 and 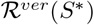 are given by Eq. [B.16], the viral strains with more lytic strategies (i.e., 0 ≤ p_m_ < 1 and γ_min_ < γ_m_ ≤ γ_max_) cannot invade the stable resident endemic equilibrium*.
ii. *If the virus-free equilibrium of model [A.1] satisfies 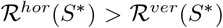, where 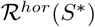 and 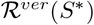 are given by Eq. [B.16], the viral strains with more lytic strategies (i.e., 0 ≤ p_m_ < 1 and γ_min_ < γ_m_ ≤ γ_max_) can invade the stable resident endemic equilibrium if 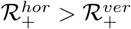, where 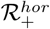 and 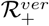 are given by Eq. [D.31]. In contrast, the viral strains with more lytic strategies cannot invade the stable resident endemic equilibrium if 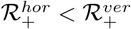*.
iii. *If the virus-free equilibrium of model [A.2] satisfies 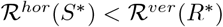, where 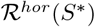 and R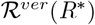 are given by Eq. [B.22], the viral strains with more lytic strategies (i.e., 0 ≤ p_m_ < 1 and γ_min_ < γ_m_ ≤ γ_max_) cannot invade the stable resident endemic equilibrium*.
iv. *If the virus-free equilibrium of model [A.2] satisfies R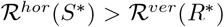, where 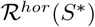 and 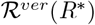 are given by Eq. [B.22], the viral strains with more lytic strategies (i.e., 0 ≤ p*_*m*_ *< 1 and γ*_*min*_ *< γ*_*m*_ *≤ γ*_*max*_) *can invade the stable resident endemic equilibrium if 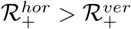, where 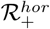 and 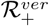 are given by Eq. [D.35]. In contrast, the viral strains with more lytic strategies cannot invade the stable resident endemic equilibrium if 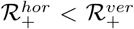*.

By theorem (1), the maximal strategies, i.e., the strategies that maximize 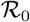, are either purely lytic strategy or the purely lysogenic strategy. For model [A.1] and model [A.2], when the resident strains are the maximal strategies, we provide remark (8).

**Remark 8** *(i) If the purely lysogenic strategy maximizes* 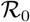, *then no other strategy can invade the endemic equilibrium for the purely lysogenic strategy*.

*(ii) If the purely lytic strategy maximizes* 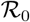, *then no other strategy can invade the endemic equilibrium for the purely lytic strategy, provided* 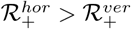 *holds*. *Otherwise, if* 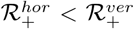, *the endemic equilibrium for the purely lytic strategy can be invaded by viral strains with more temperate strategies (i.e., p_r_ < p_m_ ≤* 1 *and γ_min_ ≤ γ_m_ < γ_r_)*.

#### D.3 Invasion of temperate phage with varying degrees of super-infection immunity

In this section, we detail the invasion analysis for model [A.2] that include super-infection. In order to control the degree of super-infection immunity, we introduce a new parameter *ϵ*, where 0 ≤ *ϵ* ≤ 1. When *ϵ* = 1, lysogens have full super immunity, viruses are absorbed into all the cells but only actively infect susceptible cells (i.e., super-infection is a ‘sink’ for viruses). When *ϵ* = 0, purely lytic viruses are absorbed into all cells and can switch a lysogen into an actively infected cell. We consider following mutant-resident system of model [A.2] with super-infection immunity mechanism (assuming resident strain is purely lytic viruses *p* = 0, mutant strain is temperate viruses with *p*_*m*_ > 0)

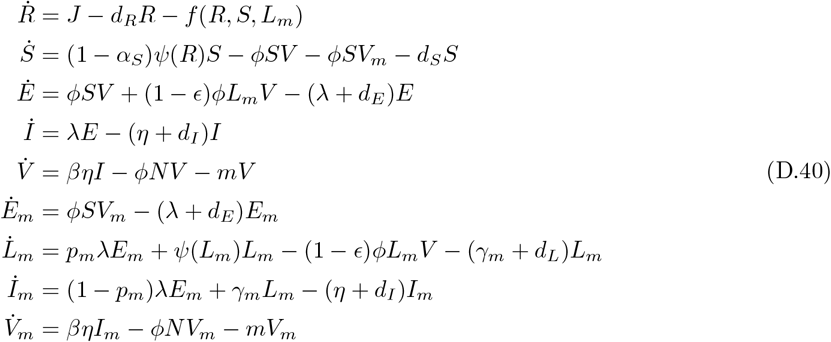

where *f* (*R, S, L_m_*) = *eψ*(*R*) (*L*_*m*_ + (1 *− α_S_*)*S*) denotes the cumulative uptake of nutrients by all cells and *N* = *S* + *ϵ* + *I* + *E*_*m*_ + *L*_*m*_ + *I*_*m*_ is the total population of hosts. The next-generation matrix of a mutant strain with viral strategy (*p*_*m*_, *γ*_*m*_) is

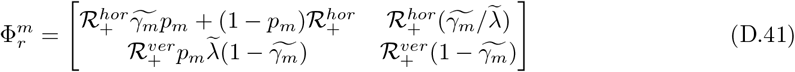

where 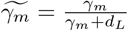, 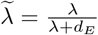 and

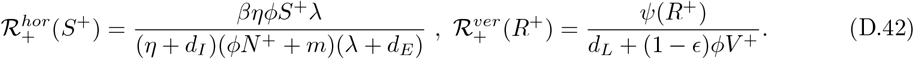

The mutant invasion fitness 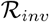 is the largest eigenvalue of NGM 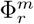 presented in Eq. [D.41].

### E Model Parameters

Here we present the parameters used to support the figures presented in this study.

**Table 1:**
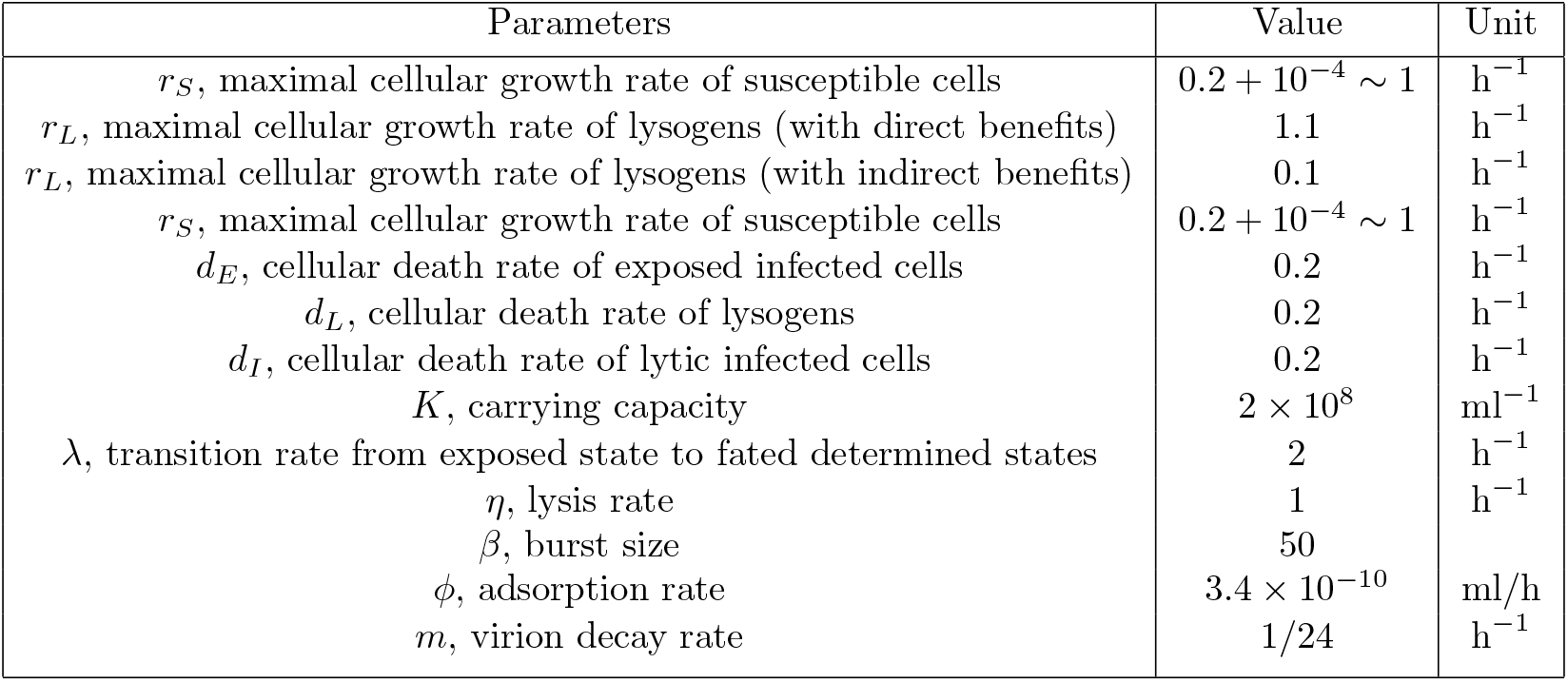
Parameters of model (A.1), source from ([6], [7])

| Parameters | Value | Unit |
| --- | --- | --- |
| $r_S$, maximal cellular growth rate of susceptible cells | $0.2 + 10^{-4} \sim 1$ | $\text{h}^{-1}$ |
| $r_L$, maximal cellular growth rate of lysogens (with direct benefits) | 1.1 | $\text{h}^{-1}$ |
| $r_L$, maximal cellular growth rate of lysogens (with indirect benefits) | 0.1 | $\text{h}^{-1}$ |
| $r_S$, maximal cellular growth rate of susceptible cells | $0.2 + 10^{-4} \sim 1$ | $\text{h}^{-1}$ |
| $d_E$, cellular death rate of exposed infected cells | 0.2 | $\text{h}^{-1}$ |
| $d_L$, cellular death rate of lysogens | 0.2 | $\text{h}^{-1}$ |
| $d_I$, cellular death rate of lytic infected cells | 0.2 | $\text{h}^{-1}$ |
| $K$, carrying capacity | $2 \times 10^8$ | $\text{ml}^{-1}$ |
| $\lambda$, transition rate from exposed state to fated determined states | 2 | $\text{h}^{-1}$ |
| $\eta$, lysis rate | 1 | $\text{h}^{-1}$ |
| $\beta$, burst size | 50 | |
| $\phi$, adsorption rate | $3.4 \times 10^{-10}$ | $\text{ml/h}$ |
| $m$, virion decay rate | 1/24 | $\text{h}^{-1}$ |

**Table 2:**
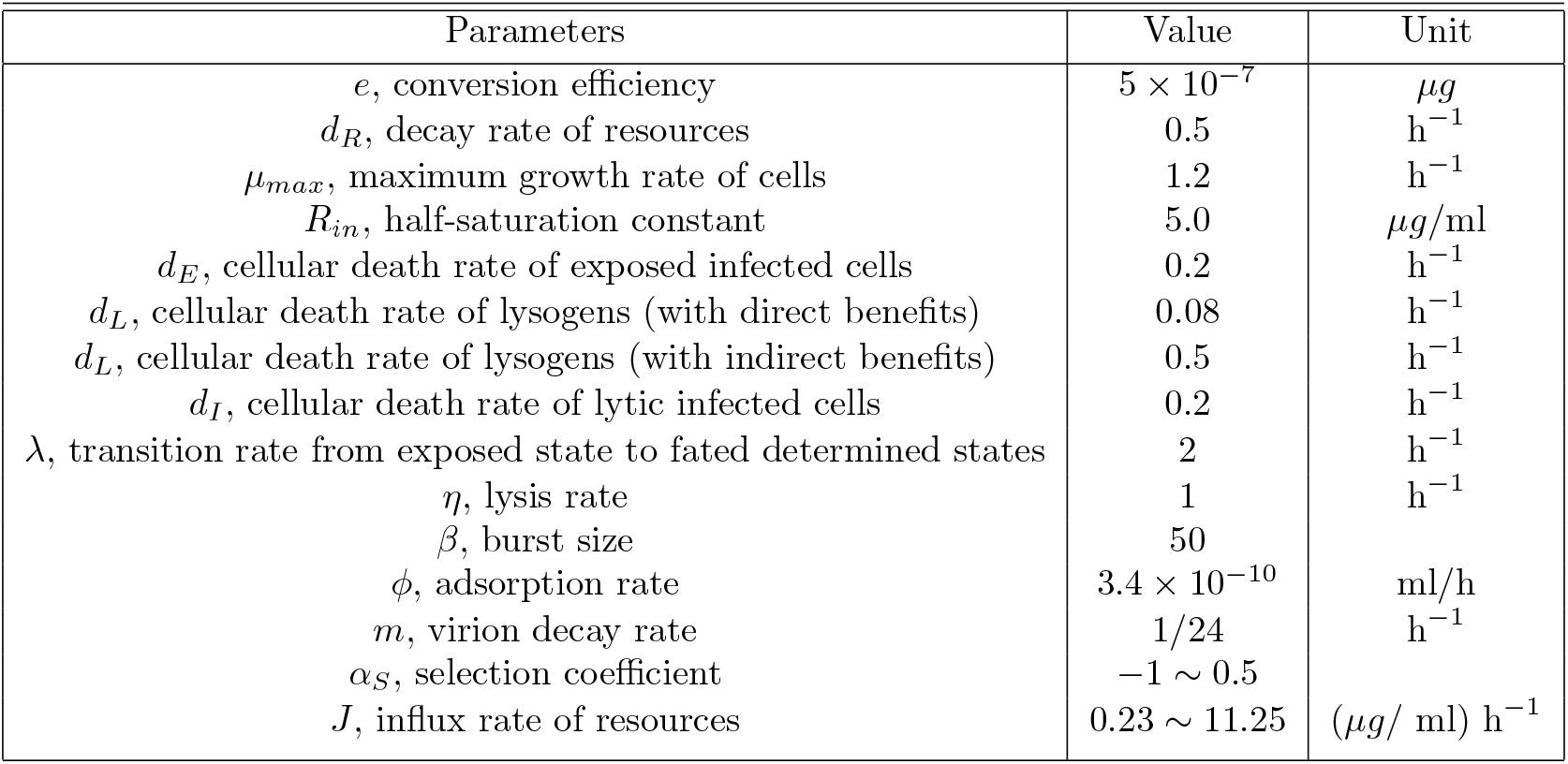
Parameters of model (A.2), source from ([2], [7])

| Parameters | Value | Unit |
| --- | --- | --- |
| $e$, conversion efficiency | $5 \times 10^{-7}$ | $\mu\text{g}$ |
| $d_R$, decay rate of resources | 0.5 | $\text{h}^{-1}$ |
| $\mu_{max}$, maximum growth rate of cells | 1.2 | $\text{h}^{-1}$ |
| $R_{in}$, half-saturation constant | 5.0 | $\mu\text{g/ml}$ |
| $d_E$, cellular death rate of exposed infected cells | 0.2 | $\text{h}^{-1}$ |
| $d_L$, cellular death rate of lysogens (with direct benefits) | 0.08 | $\text{h}^{-1}$ |
| $d_L$, cellular death rate of lysogens (with indirect benefits) | 0.5 | $\text{h}^{-1}$ |
| $d_I$, cellular death rate of lytic infected cells | 0.2 | $\text{h}^{-1}$ |
| $\lambda$, transition rate from exposed state to fated determined states | 2 | $\text{h}^{-1}$ |
| $\eta$, lysis rate | 1 | $\text{h}^{-1}$ |
| $\beta$, burst size | 50 | |
| $\phi$, adsorption rate | $3.4 \times 10^{-10}$ | $\text{ml/h}$ |
| $m$, virion decay rate | 1/24 | $\text{h}^{-1}$ |
| $\alpha_S$, selection coefficient | $-1 \sim 0.5$ | |
| $J$, influx rate of resources | $0.23 \sim 11.25$ | $(\mu\text{g}/\text{ml}) \text{h}^{-1}$ |

**Table 3:** Parameters of model (A.3), source from ([1])

| Parameters | Value | Unit |
| --- | --- | --- |
| $r_L$, growth rate of infected cells (with direct benefits) | 3.0 | $\text{h}^{-1}$ |
| $r_L$, growth rate of infected cells (with indirect benefits) | 0.2 | $\text{h}^{-1}$ |
| $r_S$, growth rate of susceptible cells | $0.75 + 10^{-4} \sim 2.25$ | $\text{h}^{-1}$ |
| $K$, carrying capacity of bacteria | $10^9$ | $\text{cells/ml}$ |
| $\delta$, fidelity of vertical transmission | 1 | |
| $\beta$, burst size | 200 | $\text{phage/cell}$ |
| $m$, dilution rate | 0.75 | $\text{h}^{-1}$ |
| $b$, probability of fusion after adsorption | $10^{-2}$ | |
| $\phi$, adsorption constant | $10^{-8}$ | $\text{h}^{-1}\text{cell}^{-1}$ |

**Table 4:** Parameters of model (A.4), source from ([2])

| Parameters | Value | Unit |
| --- | --- | --- |
| $e$, conversion efficiency | $5 \times 10^{-7}$ | $\mu g/\text{cell}$ |
| $\rho$, flow rate | 0.2 | $\text{h}^{-1}$ |
| $b$, maximum growth rate of cells | 0.7 | $\text{h}^{-1}$ |
| $R_{in}$, half-saturation constant | 4.0 | $\mu g/\text{ml}$ |
| $\nu$, prophage loss rate (with direct benefits) | $10^{-3}$ | $(1/\text{cell}) \text{ h}^{-1}$ |
| $\nu$, prophage loss rate (with indirect benefits) | 1 | $(1/\text{cell}) \text{ h}^{-1}$ |
| $\beta$, burst size | 100 | phage/cell |
| $\phi$, phage adsorption rate | $10^{-9}$ | $(\text{ml}/\text{phage}) \text{ h}^{-1}$ |
| $\alpha_S$, selection coefficient | $0.1 \sim 0.61$ | |
| $C$, resource concentration in the reservoir | $0.21 \sim 56.68$ | $\mu g/\text{ml}$ |

